# A (Sub)field Guide to Quality Control in Hippocampal Subfield Segmentation on High-resolution T_2_-weighted MRI

**DOI:** 10.1101/2023.11.29.568895

**Authors:** K.L. Canada, N. Mazloum-Farzaghi, G. Rådman, J.N. Adams, A. Bakker, H. Baumeister, D. Berron, M. Bocchetta, V. Carr, M.A. Dalton, R. de Flores, A. Keresztes, R. La Joie, S.G. Mueller, N. Raz, T. Santini, T. Shaw, C.E.L. Stark, T.T. Tran, L. Wang, L.E.M. Wisse, A. Wuestefeld, P.A. Yushkevich, R.K. Olsen, A.M. Daugherty, the Hippocampal Subfields Group

## Abstract

Inquiries into properties of brain structure and function have progressed due to developments in magnetic resonance imaging (MRI). To sustain progress in investigating and quantifying neuroanatomical details *in vivo*, the reliability and validity of brain measurements are paramount. Quality control (QC) is a set of procedures for mitigating errors and ensuring the validity and reliability of brain measurements. Despite its importance, there is little guidance on best QC practices and reporting procedures. The study of hippocampal subfields *in vivo* is a critical case for QC because of their small size, inter-dependent boundary definitions, and common artifacts in the MRI data used for subfield measurements. We addressed this gap by surveying the broader scientific community studying hippocampal subfields on their views and approaches to QC. We received responses from 37 investigators spanning 10 countries, covering different career stages, and studying both healthy and pathological development and aging. In this sample, 81% of researchers considered QC to be very important or important, and 19% viewed it as fairly important. Despite this, only 46% of researchers reported on their QC processes in prior publications. In many instances, lack of reporting appeared due to ambiguous guidance on relevant details and guidance for reporting, rather than absence of QC. Here, we provide recommendations for correcting errors to maximize reliability and minimize bias. We also summarize threats to segmentation accuracy, review common QC methods, and make recommendations for best practices and reporting in publications. Implementing the recommended QC practices will collectively improve inferences to the larger population, as well as have implications for clinical practice and public health.

## Introduction

Continuing developments in magnetic resonance imaging (MRI) have enabled progressively deepening inquiries into properties of brain structure and function. This progress has in part enabled the development of well-defined and anatomically grounded segmentation protocols for various neuroanatomical regions that can be visualized on MR images. To sustain progress in investigating and quantifying neuroanatomical details *in vivo*, the reliability and validity of brain measurements are paramount.

Hippocampal subfields are small regions, sometimes less than a millimeter in thickness, that are defined by contiguous boundaries and are distinct in their cytoarchitecture, neurochemistry, and function (Duvernoy, 2005; Insausti & Amaral, 2012). In the context of hippocampal subfields, valid *in vivo* structural measurements start with the acquisition of appropriate MR images (i.e., high-resolution T_2_-weighted images, see http://www.hippocampalsubfields.com/people/acquisition-working-group and Yushkevich et al., 2015a for details), which are segmented and labeled based on anatomical atlases developed to reflect underlying cytoarchitecture. The HSG (hippocampalsubfields.com) was developed in 2013 with the aim of developing a harmonized protocol for the segmentation of hippocampal subfields for high-resolution T_2_-weighted MRI data (Olsen et al., 2019). In our prior publications we have reviewed common imaging methods and recommended best practices for MRI protocol design for measuring hippocampal subfields *in vivo* (Wisse et al., 2020). In that prior work, we emphasized that reliable application of boundary definitions is needed to maintain confidence in the hippocampal measurement results. Although the problems arising from variations in scan quality and segmentation accuracy are not unique to hippocampal subfields, because of the small size of the targets and different application of labels along the anterior-posterior axis, the consequence of measurement error is disproportionately high. Therefore, consistent applications of quality control (QC) of collected scans and segmentation accuracy (i.e., detecting deviations in labeling of regions from the defined protocol), are important for ensuring reliable and valid measurement of the hippocampal subfields.

Although most brain MRI studies report using some form of QC, and despite occasional calls for its standardization (e.g., Backhausen et al., 2021), there are limited QC guidelines that are clearly recommended in the literature, especially in relation to specific and widely studied anatomic structures such as the hippocampus. When testing hypotheses involving MRI-derived measurements, QC provides a means to mitigate measurement error that can lead to Type I and Type II errors, and subsequently improves the reproducibility of results (Elyounssi et al., 2023). Therefore, reporting QC details of the segmentations are necessary to support interpretation of hippocampal subfield measures correlated with function and cognition across the human lifespan, and potential application as biomarkers of disease processes.

## Survey on QC

Currently neither concrete recommendations for best practices of QC for hippocampal subfield measurement nor minimum reporting standards exist in the literature. We recently began to address this gap by surveying the views and approaches to MRI QC from the broader community of Hippocampal Subfields Group (HSG) investigators who work with measures of hippocampal subfield structure. Thus, as investigators concerned with assessing the role of hippocampal subfields in development, aging, cognition, and neuropathology, our goal is to provide a guide to QC that will be effective in this specific application to subfield segmentation.

An extensive survey assessed the views and approaches to QC of hippocampal subfield segmentation in the HSG community and was distributed using the HSG’s listserv, social media, and website. Survey responses were collected from 07/11/2022 to 10/14/2022. The survey was completed by 37 respondents each representing a different laboratory spanning 10 different countries and 4 continents. Respondents’ research represented the study of hippocampal subfields across the lifespan (from 0 years - 75+ years) and in healthy and diseased populations (see Supplementary Material for detailed breakdown of respondent demographics).

Of the 37 respondents, 81% considered QC to be “very important or important” and 19% considered QC to be “fairly important”. While all respondents considered QC to be important to some degree, only 46% reported on their QC practices in prior publications. This response highlights the mismatch between the importance of QC and the inconsistent standards of reporting QC in published studies. In many instances, this may not be due to absence of QC but merely ambiguous guidance on relevant details to report (see Supplementary Material for detailed survey results).

## QC Guidelines

The goal of this guide is to provide guidance covering the various decision points investigators may encounter during the QC process. We take an *à la carte* approach and anticipate an investigator may choose to implement some or all of the practices we review, and that this will vary across studies. Our intention is to provide an overview of each stage of QC to allow for the investigator to make informed decisions for their study and follow best practice recommendations on reporting the procedures they implement. The goal of this guidance is to improve the quality of data and transparency in reporting within the hippocampal subfields literature.

Although the many protocols to delineate hippocampal subfields vary meaningfully in both their label composition and defined boundaries (Yushkevich et al., 2015a), a common set of QC procedures will be applicable to any manual or automated segmentation protocol. In the following sections, we propose best practices for QC of acquired MR images and labeling to measure hippocampal subfields. We also provide concrete suggestions and illustrations of how to identify, correct, and report segmentation errors. Below, we briefly address several topics related to the QC process, such as excluding data of questionable quality, measuring the reliability of QC interventions, reducing bias in subfield measurement, and special considerations for implementing QC procedures in longitudinal studies (see schematic summary of QC procedures in Figure 1).

**Figure 1.**
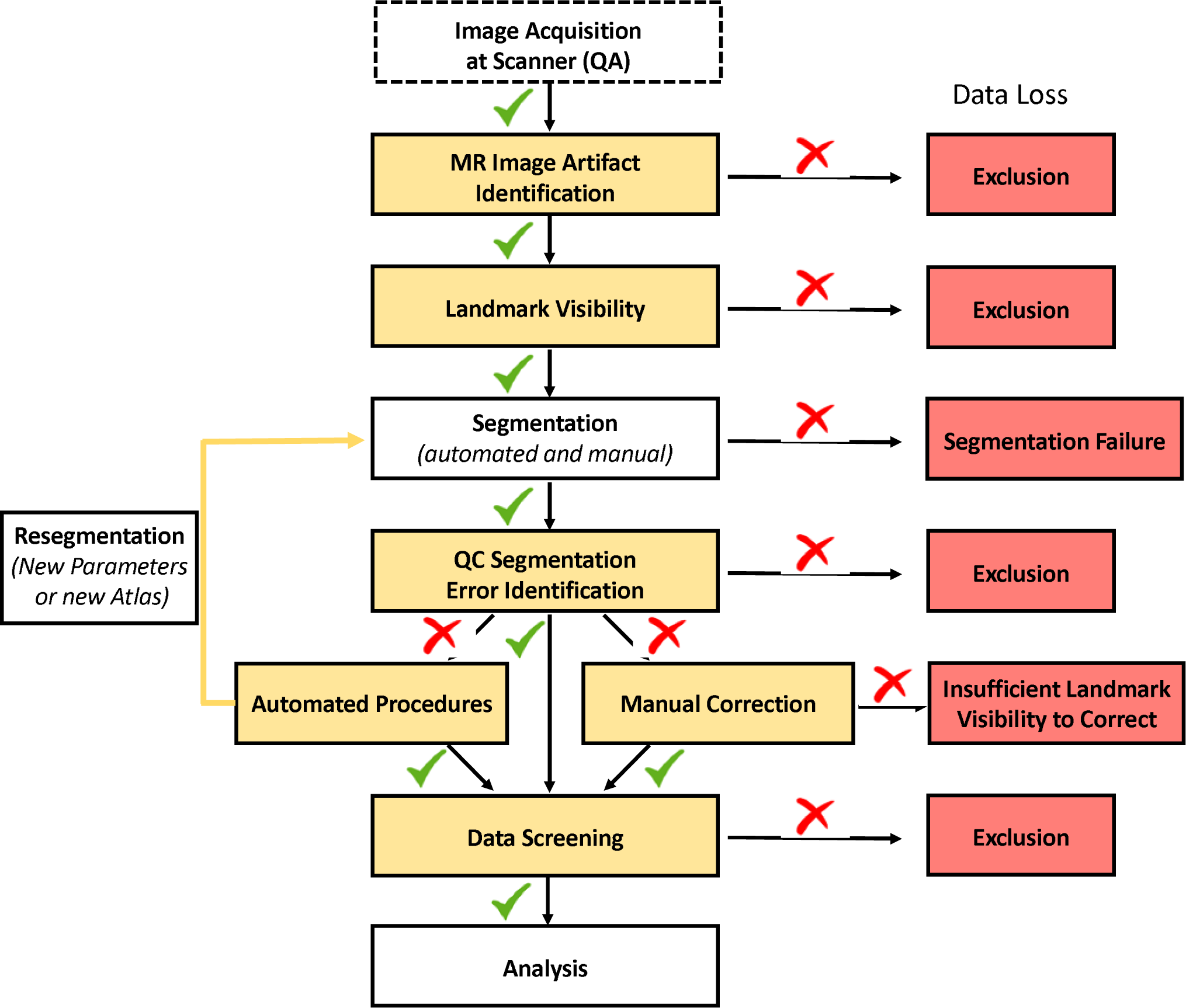
Illustration of the QC process and investigator-guided decision making for data quality. Green checkmarks indicate passed QC, while red cross marks indicate failed QC.

## QC of MR Images

Quality of MR images directly affects the reliability and validity of segmentations, and by extension, the reproducibility of results. Two related aspects of MR image QC are the identification of artifacts affecting overall image quality and image features that impede the ability to visualize neuroanatomical landmarks. Prior to segmenting hippocampal subfields (either manually or automatically), a critical first step in the QC process is assessing the quality of acquired images for imaging artifacts, poor contrast, and insufficient landmark visualization. Across the investigators surveyed by the HSG, over 89% of respondents reported conducting QC of MR images, and 94% of those exclude images due to quality issues. Segmentation of hippocampal subfields relies on the visualization of specific landmarks, typically on T_2_-weighted high-resolution images (Wisse et al., 2020), in order to determine outer boundaries of the hippocampus and the inner boundaries between subfields (see *QC of Landmark Visibility* section). Thus, our discussion and illustration of QC for MR images in the following sections are applied in the context of hippocampal subfield measurement from oblique coronal T_2_-weighted scans with high in-plane resolution and orientation roughly orthogonal to the hippocampus as recommended by the HSG (for a discussion of T_1_-weighted image QC not specific to hippocampal subfields see Alfaro-Almagro et al., 2018; Rosen et al., 2018).

### MR Image Artifact Identification

#### Description of the Problem

MR images are prone to artifacts during acquisition due to a variety of factors, including participant movement, metal implants, magnetic field inhomogeneities related to head geometry and tissue compartments, and mechanical faults in the gradient coils. These result in suboptimal image quality and the ensuing “artifacts”. These artifacts affect the quality of the data, which in turn reduces the quality of the segmentation. Although images do not need to be perfect to have valid measurements of brain structures, there is a minimum standard of data quality that often leads investigators to exclude scans as a first step in QC.

Among surveyed investigators, motion artifacts (e.g., ghosting) and susceptibility artifacts (e.g., image distortion due to metallic dental work) (Figure 2) were the most common examples of imaging artifacts flagged in QC of T_2_-weighted images. Motion artifacts can degrade the clarity of the image due to blurring of boundaries between tissue compartments and loss of image contrast (Reuter et al., 2015). Of note, the majority of respondents identified problems related to motion as the most common cause of exclusion (70%). For example, ghosting artifacts appear as rings or streaks within slices and are due to motion. Artifacts due to reconstruction errors can occur near high contrast boundaries and can also appear as multiple lines in the image that alternate between bright and dark coloring (i.e., rings or bands) (Bellon et al., 1986). Susceptibility artifacts also affect the tissue appearance on MR images as local magnetic field inhomogeneities are translated into structural distortions and signal dropout. For example, the presence of an implanted metallic object with significant magnetic susceptibility (e.g., metallic dental work) results in pronounced inhomogeneities in the B_0_ static magnetic field and subsequently issues in the reconstruction of the affected areas. These artifacts are exacerbated at higher field strengths (Dietrich et al., 2008) and independently, as well as collectively, lead to insufficient image contrast that could severely distort hippocampal subfield segmentation.

**Figure 2.**
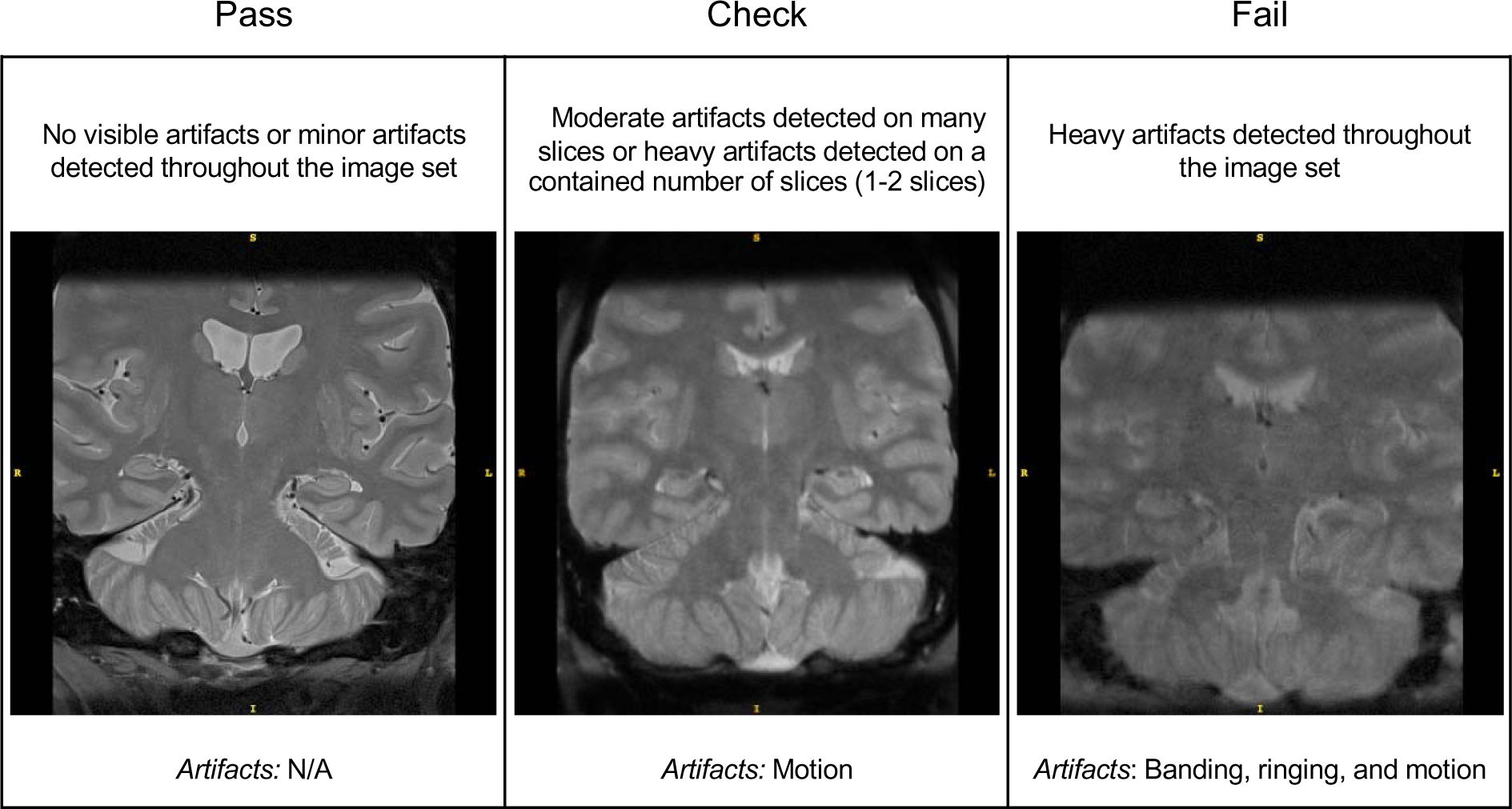
Examples of the quality of T_2_-weighted images according to rating categories of “Pass” (left panel), “Check” (middle panel), and “Fail” (right panel). In this example, image quality rated as “Pass” and “Check” are considered passable. However, those in the “Check” category may be at higher risk of subsequent segmentation errors. The types of imaging artifacts present in the example images are noted.

#### Review of Current Approaches and Recommendations

As an overall summary of image quality, a practice noted by a number of investigators surveyed is to quantify the image signal-to-noise ratio (SNR) and contrast-to-noise ratio (CNR). MR image SNR is the proportional mean signal of a region of interest to background noise (typically the standard deviation of the signal sampled from air space), whereas CNR refers to comparison of the mean signal in a region of interest to a reference region (e.g., gray matter to white matter) in proportion to background noise. Thus, SNR provides a global estimate of image quality, whereas CNR provides a local estimate of contrast between tissue types. Additional inspection of the MR image for artifacts can be performed manually or with automated tools (detailed below).

##### Manual Image Quality Screening Procedures

Qualitative review is a common approach in assessing the quality of MR images. This type of approach entails the visual inspection of all images in a data set, which is usually done by one or more trained raters. Manual QC procedures are based on investigator judgment and experience. As some degree of imaging artifact is often tolerated during segmentation, the rater decides whether artifacts are severe enough to undermine confidence in subsequent segmentation, including anticipating a critical segmentation failure with automated software. Based on the results of the inspection, low-quality images are often excluded from the data set.

Rater’s review of image quality is sometimes reported in methods sections of papers, but the specific procedure for this is rarely explicated. In some applications, the QC decision appears to be a binary choice (i.e., include or exclude), whereas in other instances, an ordinal rating scale may be used to describe the quality of the image or severity of artifacts (e.g., 0 = no artifact/pass, 1 = minimal artifact/check, 2 = severe artifact/fail). We recommend using a rating system (either a binary or multi-point ordinal scale) to determine if an image is of sufficient quality for segmentation. This approach aids investigators in specifying the operational criterion used to determine the image quality as sufficient and improves the transparency and reproducibility of the methods. For examples of scales and criteria related to imaging artifacts see Backhausen et al. (2021), Ding et al. (2019), and Rosen et al. (2018). Because of subjectivity in this procedure, describing the criteria for the decision and the reliability of the procedure (e.g., kappa statistics between- or within-rater; minimum 0.75 indicates a strong level of agreement; Fleiss et al., 2003) is important for ensuring the quality of the generated data and reproducibility of the findings.

##### Automated Image QC Methods

As manual image quality evaluation can be labor intensive and time consuming, especially when working with large data sets, automated approaches present an efficient alternative. There are several automated tools available to assess the quality of MR images (e.g., MRIQC, Esteban et al., 2017; LONI QC, Kim et al., 2019) and a semi-automated approach using machine learning (Ding et al., 2019). These tools provide internal consistency and easy-to-read outputs with user-rating options (e.g., html pages for MRIQC). Some of the tools also provide volumes of the structures in question and flag statistical outliers, which is particularly useful in dealing with very large data sets (see *Measurement Data Screening* section for additional information on statistical outliers). We recommend reporting the parameters for exclusion and the level of artifacts or image quality tolerated by the automated method (e.g., Ding et al., 2019). An optional addition for those who use a quantitative SNR or CNR measurement is to exclude scans with low ratios in the areas of interest (i.e., hippocampus) compared to the reference region, with SNR less than 40 and CNR less than 10 (Magnotta et al., 2006). A caveat to this recommendation is that the procedures were developed for the data collected in healthy participants and have not been validated in the images with significant structural pathology (e.g., hippocampal sclerosis in temporal lobe epilepsy).

Automated approaches have high internal consistency. However, it is important to note that their application to detecting poor image quality may have some disadvantages. These tools can be used with little user knowledge of relevant parameterization programmed into the method, and users may be unable to change certain steps or parameters during the automated assessment. Moreover, automated methods may not be as accurate as experienced human raters in detecting subtle distortions in images. In choosing between manual and automated procedures, the investigators can weigh their knowledge of the methods against the expertise of the team and available time.

##### One final note

During QC decisions, investigators should consider the context of the population under study. For example, individuals with more severe diseases are likely to have more artifacts on MRI. In these instances, extremely conservative QC practices may lead to disproportionately excluding persons with high disease severity, which leads to another form of bias due to underrepresentation of the population under study. Therefore, an investigator may use their knowledge of the population in evaluating the risk-benefit tradeoff of their data QC practices.

### QC of Landmark Visibility

#### Description of the Problem

Brain region segmentation in manual and automated protocols is based on anatomical landmarks that are visible on MRI and correlate with histologically identifiable macro- and micro-structural tissue features. As the correspondence of MRI labels to histology constitutes the basis of construct validity of *in vivo* measurements, any artifact or distortion that interferes with visualization of key landmarks in the segmentation protocol weakens the validity (Wisse et al., 2020). Of particular importance for hippocampal subfield segmentation is the visualization of the stratum radiatum lacunosum moleculare (SRLM), a thin, layered sub-1-mm^3^ structure (de Flores et al., 2020). It spans the anterior-posterior length of the hippocampus and forms a layer of the cornu ammonis (CA) regions and subiculum. It also serves as a critical landmark for determining the internal boundary between dentate gyrus (DG) and CA subfields, or subiculum (Duvernoy, 2005; Insausti & Amaral, 2012). In addition, the visualization of the SRLM in the hippocampal head is important for the identification of structural digitations, which play a critical role in determining the presentation of subfields in the anterior hippocampus (Adler et al., 2018). In T_2_-weighted images of sufficient quality as we presume here, the SRLM should be clearly visible across most hippocampal slices as a dark band perpendicular to the anterior-posterior axis of the hippocampus (Figure 3).

**Figure 3.**
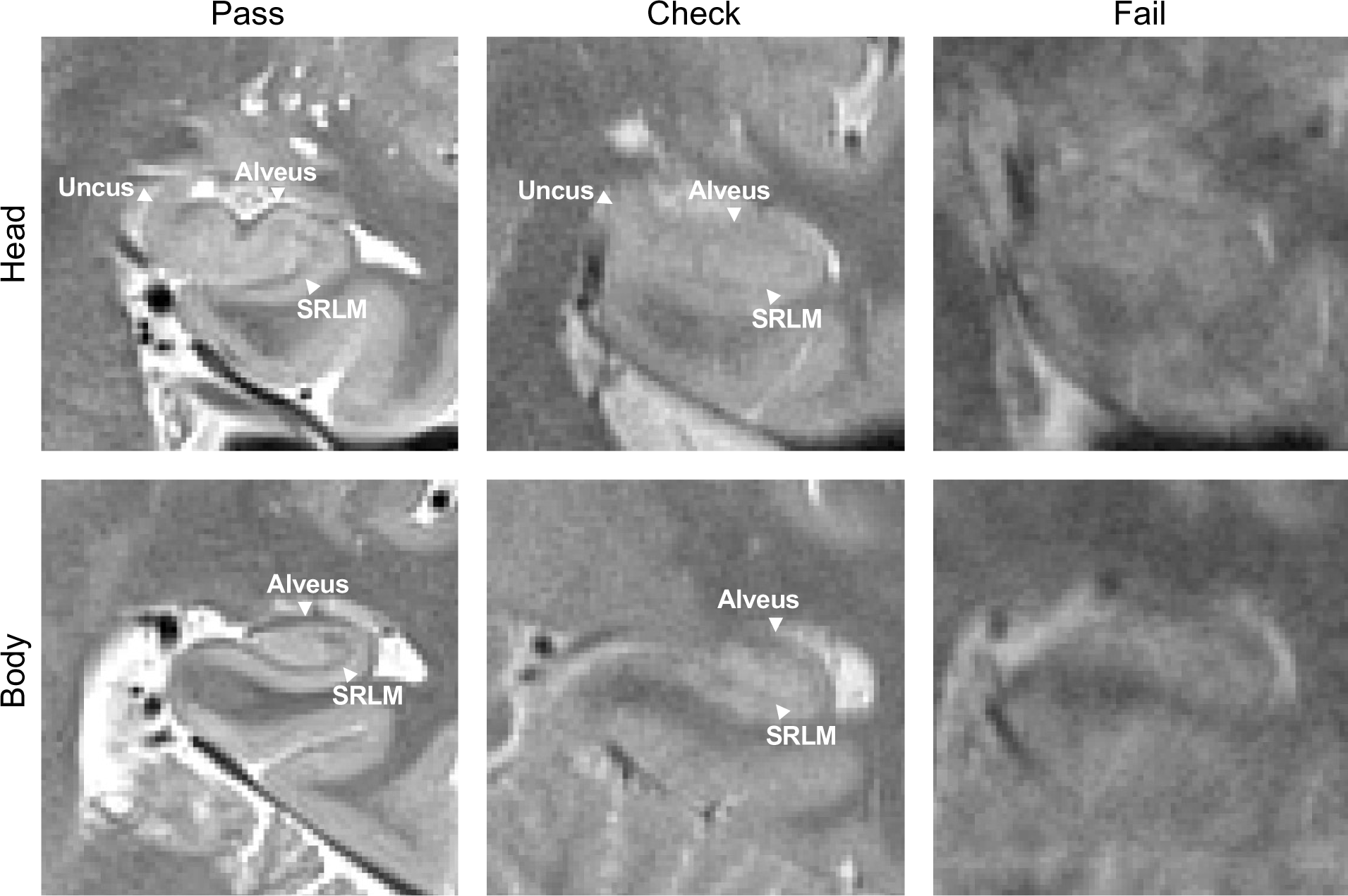
Examples of scan quality ratings based on the landmark visualization of the SRLM, a critical landmark for segmentation in the head (top panel) and body (bottom panel) of the hippocampus. Additional landmarks, the alveus and uncus, are also depicted. Note: Definitions of quality may differ between investigators but there should be consistency in the application of operational definitions.

Because of similar decision-making regarding MR image quality, the procedures for evaluating landmark visibility follow the same steps and are often performed together. Although issues of image quality and landmark visibility apply to any region of interest, segmentation of hippocampal subfields is somewhat unique in its contiguous regions that share internal boundaries. If one boundary is incorrectly determined due to poor visualization of a landmark, the error can propagate across all subfield measurements, diminishing the validity of each subsequent label. An additional nuance of hippocampal subfield segmentation is that the label definitions for a subfield often shift on the anterior-posterior axis, and in many cases the same region will have separate labels anterior-to-posterior and by hemisphere. Therefore, depending on the protocol used, QC decisions may vary across different levels of measurement. For example, QC decisions may differ by slice, subfield, subregion, or hemisphere. In addition, an investigator may evaluate landmark visibility only within the limited range of the hippocampus sampled (e.g., many subfield protocols are exclusive to the hippocampal body) or decide to use measurements from one region or hemisphere and not others due to differences in quality (see *Approaches to Data Exclusion and Recommendations* considerations of missing data). Further, QC decisions may vary depending on the imaging modality. Extensive discussion of modality considerations in QC falls outside the scope of this guide. However, to provide an example, those employing hippocampal subfield segmentations to estimate the volume of these structures rely on information across numerous slices whereas those applying segmentations as a mask in fMRI may use measurements from only a few slices.

Notably, among those survey respondents who conducted QC, the SRLM was the most frequently identified landmark reviewed in QC (42% of investigators). However, decisions regarding exclusion based on SRLM visibility varied across investigators. For example, multiple respondents noted that scans were excluded only if there were issues with SRLM visualizations on multiple consecutive slices, while others did not specify criteria but included SRLM visibility as a factor in a subjective judgment combining multiple artifacts and problems. See Figure 3 for examples of different SRLM visualization quality on T_2_-weighted images.

In addition to the SRLM, other prominent landmarks (e.g., alveus, fimbria, endfolial pathway, ambient cistern, or uncus depending on specific segmentation atlases used) should be visible in order to determine the border of contiguous hippocampal subfields and the anterior-posterior transitions from hippocampal head, body, and tail. For example, the uncal apex is often used for identifying the transition from hippocampal head to the body (Malykhin et al., 2007), which marks a change in the morphometry of the subfields for labeling. The lamina quadrigemina (LQ) and visualization of the fornix are additional landmarks used to determine the posterior boundary of the hippocampal body and transition to tail. Bender et al., (2018) noted that different ranges can be established by hemisphere so long as at least one of the four colliculi of the LQ is visible. The fornix is also used in some protocols to define the most posterior slice of the hippocampal body, namely the slice before the fornix is fully visible or clearly separates from the wall of the ventricle (Malykhin et al., 2007, 2010).

Apart from landmarks used to identify anterior and posterior portions of the hippocampus, structures such as the alveus and fimbria facilitate the identification of outer boundaries to exclude external white matter and partial voluming with cerebrospinal fluid and choroid plexus in the gray matter labels. The alveus is a thin white matter structure enveloping the dorsal aspect of the hippocampus. It appears as a dark band on T_2_-weighted images on the dorsal edge of the hippocampus and is contiguous with the fimbria in the posterior hippocampus. This structure helps identify the boundary of the hippocampus and the shape of digitations in hippocampal head and can aid in identifying the first slice of the hippocampus as it distinguishes from the amygdala. Visualized external white matter structures often serve as a landmark to identify the superior boundary of the CA regions throughout the length of the hippocampus. In the posterior hippocampus near the tail region, the fimbria is continuous with the columns of fornix that form a sulcus at the junction with the DG. In addition, it serves as a posterior landmark to the hippocampus (for depiction of selected landmarks see Figure 3). While the specific landmarks referenced may differ between protocols, clear visualization of these landmarks is essential to reliably identify the outer boundaries of the hippocampus and its inner subfield boundaries.

#### Review of Current Approaches and Recommendations

To our knowledge, there are no automated methods for evaluating landmark visualization independent of general image quality and artifacts. Therefore, these evaluations are performed manually and completed using software such as ITK-SNAP (www.itksnap.org; Yushkevich et al., 2006), FreeSurfer’s FreeView application (Fischl et al., 2002), FSLeyes (McCarthy, 2023), and Analyze (AnalyzeDirect, Overland Park, KS, USA) to view the slice images. Common practice requires that raters be familiar with neuroanatomy on MR images in reference to the protocol used to make sound judgments about landmark visualization.

Following the recommended practices for QC of MR image artifacts mentioned in the prior section, we recommend using rating scales to determine the quality of landmark visualization required for hippocampal subfield segmentation. For example, on a 3-point scale, scans could be identified as “Pass/Clearly Visible”, “Check/Somewhat Visible”, or “Fail/Not Visible” based upon the visibility of the selected landmark (e.g., SRLM). This or other similar rating systems could be used to make determinations of exclusion across the multiple criteria we have discussed. As noted above, QC decisions may vary across different levels of measurement by slice, by region, or by subfield and hemisphere.

As with artifacts, the rating procedure for landmarks is somewhat subjective, and consistency in the decision within a research team applied to a data set should be prioritized. To allow for the transparency necessary for replication across research groups and references in the extant literature, we recommend that investigators describe the rating procedure, including the specific landmarks examined for the chosen segmentation atlas and criterion used to determine the rating system. Further, investigators should report reliability of the raters in the decision (e.g., a kappa statistic) to demonstrate consistency in the subjective decisions made using the defined criteria. This will allow some continuity of methods in the literature, even if different investigators refer to different criteria based on the study sample (e.g., healthy vs. patients), segmentation protocol, or imaging modality. When trained raters cannot clearly identify the landmarks, the scan is judged to be of insufficient quality and is typically excluded from measurement.

Finally, like the QC of MR image quality, QC of landmark visibility should consider the context of the population under study. For example, in the context of disease, the SRLM might be more difficult to visualize with increasing severity.

## QC of Hippocampal Subfield Segmentations

### Description of the Problem

The procedure for determining the accuracy of hippocampal subfield segmentation labels depends on whether the segmentation was generated using manual or automated methods. Among the survey respondents, 88% report having employed automated segmentation methods and 49% manual segmentation methods for delineating hippocampal subfields. Many investigators applied either of these methods depending on the data set.

When using manual segmentation, rater reliability should be established before segmentation commences. Besides being an indicator for accurate segmentation, a high rater reliability of a manual segmentation protocol also implies accurate identification of severe segmentation errors that deviate from the protocol during QC. Of the respondents who used a manual approach for segmenting hippocampal subfields, 77% reported assessing inter- or intra-rater reliability of the protocol. Whereas QC of the labels may be done concurrently with manual segmentation, labels from automated segmentation must be vetted afterward. Additionally for automated segmentation, reliability between the manual and automated segmentation should be confirmed (e.g., Yushkevich et al., 2010). Even though automated segmentation has high reliability (Bender et al., 2018), it can produce errors with high consistency.

### QC Segmentation Error Identification

Among investigators surveyed, 95% reported reviewing the quality of subfield segmentations, with 63% providing specific examples of errors identified. The survey response notably highlights that, despite differences in existing protocols, there are some common types of segmentation errors: 36% reported issues related to overestimation (e.g., inclusion of choroid plexus or cerebrospinal fluid, overestimation errors due to partial volume effects or voxels erroneously extending into the fimbria) or groups of pixels labeled unattached to the hippocampus; 13% reported mislabeling of hippocampal subfields (e.g., labels overextending internal boundaries, or sulcal cavities labeled as tissue); and 9% reported underestimation of regions (e.g., labels not fully extending to the boundary, or groups of pixels dropped from within the label). Errors in segmentation labels may result from multiple sources: some related to image quality and landmark visualization that we have already discussed, and others reflecting specific properties of the automated software (for examples see Wang & Yushkevich, 2013). It is important to note that in applying automated software, the segmentation atlases validated in one data set can produce segmentation errors when applied to new data sets. Bias in the frequency or type of errors may also differ depending on the population under study—e.g., certain errors may be more common in particular populations (patients, childhood development) as compared to healthy adults (see Breakout Box 1). Identifying such errors using QC allows for the efficiency of an automated approach while ensuring measurement validity.

#### Review of Current Approaches and Recommendations

Despite the similarity among segmentation error types, there was little consensus among the survey responders on the steps taken to identify these errors. Further, these steps differed between those investigators who performed manual as compared to automatic segmentation procedures. In both automated and manual segmentation, QC for segmentation errors depends on knowledge of the standard anatomy and the segmentation atlas or protocol used to define hippocampal subfields. Our key recommended practice is to clearly describe in the methods section of a paper how segmentation labels were reviewed for errors.

For manual procedures, labels are commonly reviewed during segmentation as well as post-segmentation. Expert knowledge of the trained, reliable rater is typically depended upon for identifying errors. We recommend investigators report rater reliability for the manual segmentation protocol, with intra-class correlation assuming random raters (ICC[2,1]; Shrout & Fleiss, 1979) if measuring volumes, or report dice coefficients if using the segmentations as masks on other imaging modalities. Investigators should also report if segmentations have been reviewed by trained independent raters. For reliability assessment that enables the user to partition multiple sources of unreliability, see Brandmaier et al. (2018).

For automated segmentation procedures, the most common approach to identifying errors is visual inspection of the output using specialized software to open segmentation files (e.g., ITK-SNAP, www.itksnap.org; Yushkevich et al., 2006). For those respondents on the survey who indicated visually inspecting automated segmentations, 56% sought large or obvious errors. If an investigator is considering manual correction of automated segmentation errors (e.g., semi-automated methods), we recommend using segmentation quality ratings to correct only the most severe errors, reducing burden as well as possible introduction of human error in the process. Scales should have operational criteria to define error severity levels and establish reliability of the rater(s) (e.g., kappa statistics for within- or between-rater reliability). Criteria across protocols do not need to be identical. Instead, the best practice is to provide operational definitions that allow investigators to consistently identify errors that threaten validity. For example, investigators may define error severity based on the percentage of the label affected (see Figure 4 for an example of an error rating scale). Another approach has been to determine the extent of subfield labels affected on multiple slices along the longitudinal axis of the hippocampus as an index of severity (see Figure 5 for an example). In this approach, only errors affecting a certain length (e.g., 3-7 mm, 3 or more slices) of the longitudinal axis were considered severe and subsequently corrected or excluded. There is also value added by concretely defining specific errors that commonly occur (see Figure 4) to aid rater training and promote consistency within and between raters.

**Figure 4.**
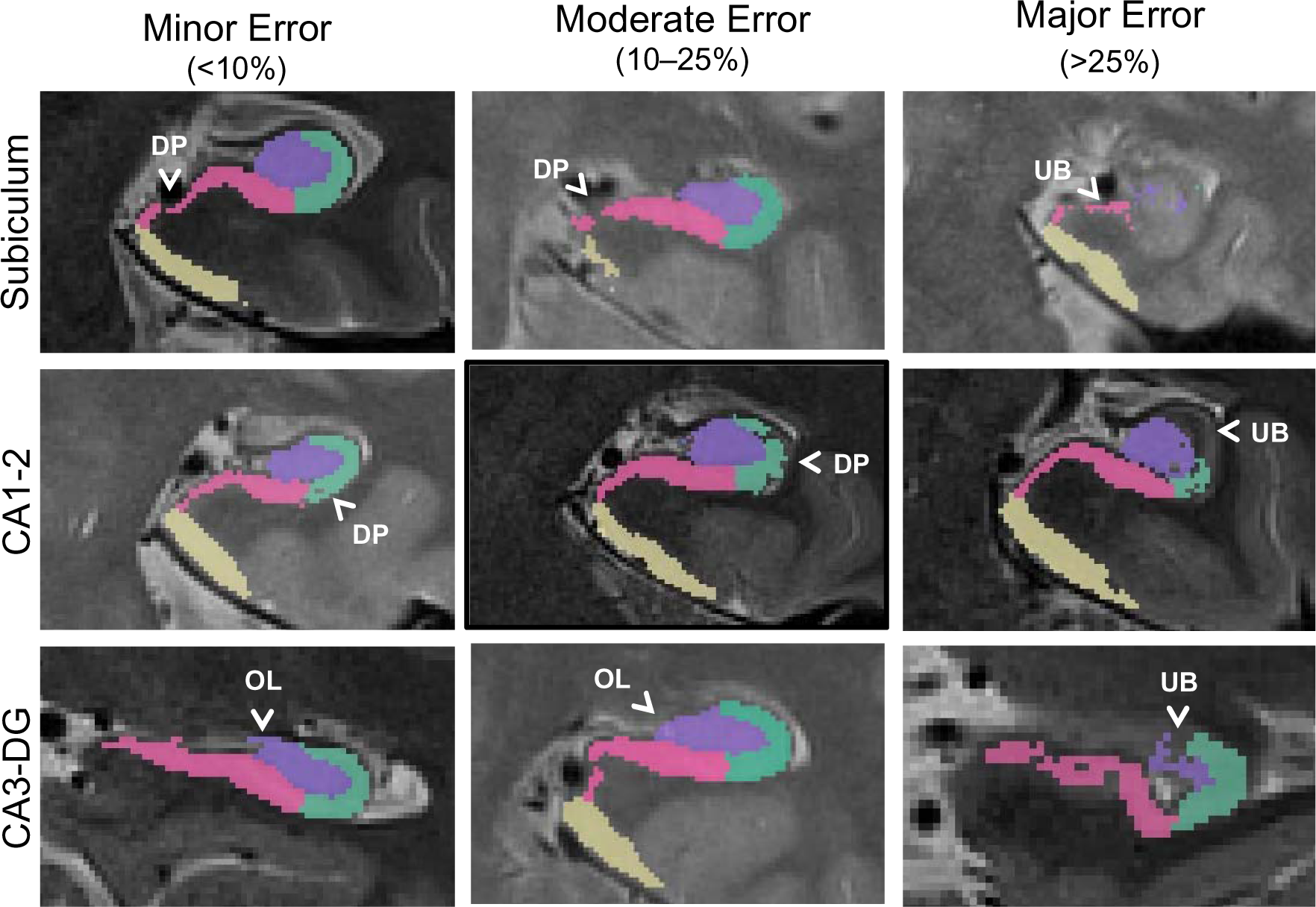
Example QC approach using a 4-point error severity scale from a published and validated protocol (Canada et al., 2023). In this example protocol, errors could be 0- not present (not pictured here), 1- minor (<10% of label affected), 2- moderate (10–25% of label affected), or 3- major (>25% label affected) and were categorized by type. Only major errors are corrected in this protocol to mitigate human bias. Overestimated label (OL), underestimated boundary (UB), and dropped pixel (DP) errors are depicted.

**Figure 5.**
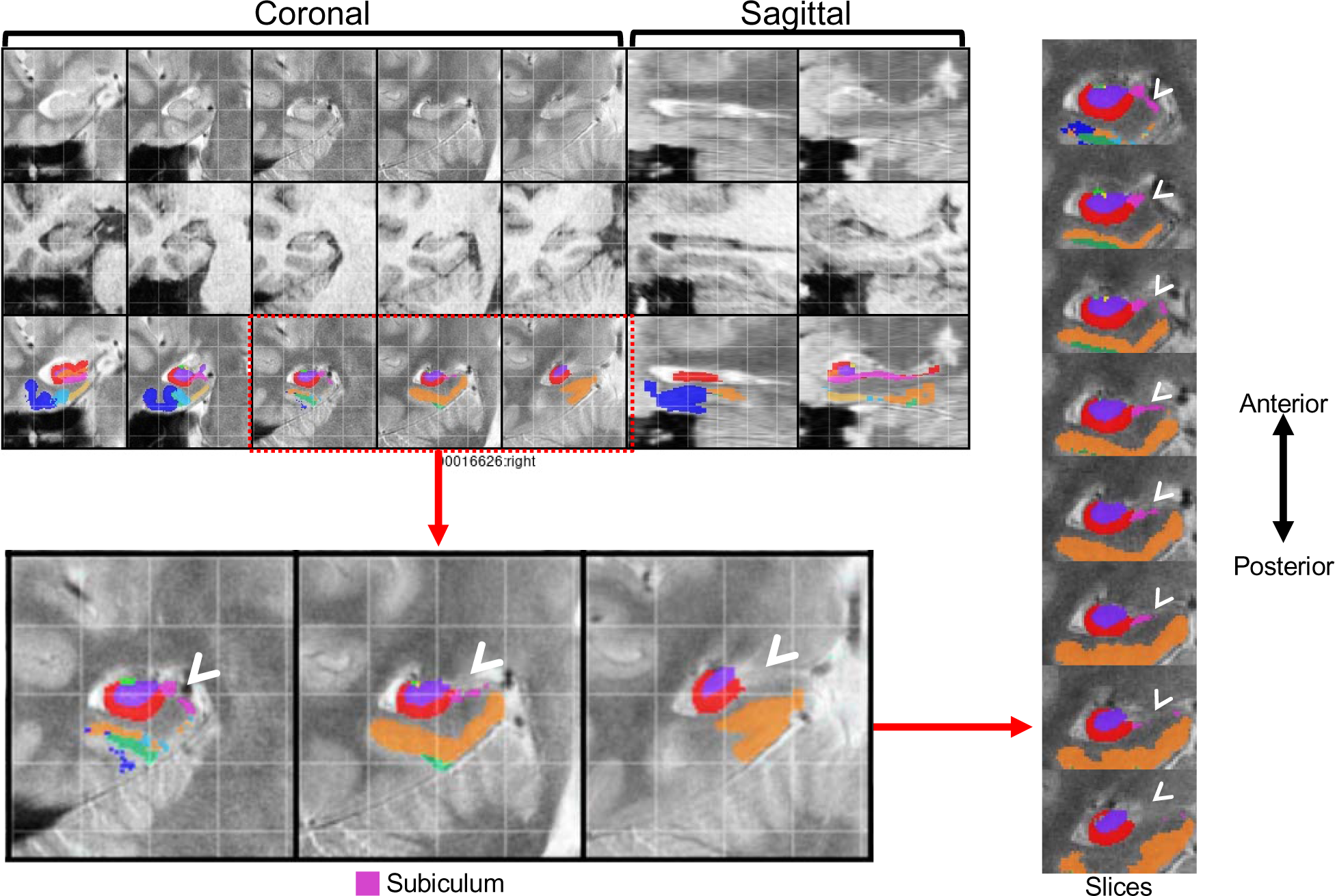
Example QC approach from https://www.youtube.com/watch?v=XHXu-AGR6pE for the PMC segmentation atlas (Yushkevich et al., 2015b) applied to data collected using the parameters reported in Daugherty et al. (2016). In this approach, QC mosaic screenshots generated by ASHS are examined by hemisphere to detect potential errors for each subject. Segmentations are subsequently fully reviewed slice-by-slice only for regions with identified errors. This protocol recommends correcting only larger errors that are present on a predetermined number of slices (3 slices is used in the linked demonstration). In the depicted example, major errors were present in the right hemisphere QC screenshot for the subiculum (pink) label. In the full review of the segmentation, errors in the right subiculum label were present in 8 slices. Following this QC approach, the depicted subject would require manual correction of these errors or be excluded from analyses.

The main purpose of identifying errors on a severity scale is not to have perfect segmentations but rather to identify cases with egregious threats to validity. Multiple investigators reported using a 4-point scale to quantify the severity of segmentation errors as not present (0), minor (1), moderate (2), or major (3), while others reported using pass/fail ratings. In addition, 31% of respondents reported examining inter- or intra-rater reliability of error identification ratings. To determine severity of errors, some investigators employ a group review where either the segmentation or screenshots are examined by multiple people simultaneously and decisions on the presence or absence of an error are determined by group consensus. Reporting on the approach(es) that investigators take to identify segmentation errors, 10% of respondents stated that segmentation error identification was always completed by multiple raters, 60% mentioned that errors were sometimes reviewed by more than one rater, and 30% said that only a single rater reviewed errors. It is important to note that the variability in the number of raters that review segmentation errors across labs may be due to differences in the availability of personnel necessary to review segmentation quality. It is important to note that in cases involving very large data sets (e.g., Alzheimer’s Disease Neuroimaging Initiative (ADNI) data sets), it may not be feasible to QC all of the collected scans for segmentation errors. As such, at least a subset of the data set should be QC-ed in order to get a sense of the data quality.

Similar to the judgments of MR image quality, the manual evaluation of segmentation errors is also subjective; thus, it requires standardized approaches to promote consistent decisions. There are several resources available to provide some options for standardized protocols: 1) MAGeT-related QC (https://github.com/CoBrALab/documentation/wiki/MAGeT-Brain-Quality-Control-(QC)-Guide), 2) HippUnfold’s automated QC (DSC overlap with a deformable registration), and 3) MRIQC (Esteban et al., 2017). Additionally, the examples of manual evaluations from Canada et al. (2023) and Wisse (https://www.youtube.com/watch?v=XHXu-AGR6pE) demonstrate that investigators can use different criteria but similarly implement the recommended best practice to identify severe segmentation errors. Using existing QC protocols as a resource provides operational definitions and criterion that can be applied or modified for an investigator’s particular data set.

The inability to confidently appraise or correct segmentations due to image quality can be a related, but distinct, problem that also can lead to exclusion of images. However, if an image has passed the QC for artifacts and landmark visualization, this issue is less likely to occur (see MR Image Quality).

### Resegmentation, Manual Correction, or Exclusion of Cases with Errors from Automated Segmentation

#### Description of the Problem

Segmentation errors often affect multiple subfields because they share boundaries. Following QC of images and segmentations, decisions for resegmentation, correction, or exclusion may differ by subregion or hemisphere. As hippocampal subfields are part of a whole, the choice to exclude any single subfield label on a particular slice or for a particular participant would preclude interpretation of generalized hippocampal effects.

Because of these challenges to hippocampal subfield segmentation, we review multiple approaches to support measurement validity after identifying automated segmentation errors: manual correction, submitting images for re-segmentation by automated methods, and data exclusion. Each approach calls for different degrees of rater expertise and time investment. The choice to exclude cases versus correcting errors can depend on several factors. Among survey respondents who reported either practice, 25% of investigators made the decision to correct or exclude based on the sample size available (e.g., excluding subjects in large studies versus correcting segmentations in smaller studies), 31% made their decisions based on the uniqueness of the population (e.g., retaining rare cases), 31% made their decisions based on bias in segmentation errors (e.g., error frequency related to certain demographics; responses not mutually-exclusive). However, 49% of survey respondents who correct or exclude segmentations reported that they did not consider sample or study specific factors in decisions.

We describe common procedures for each approach and a few considerations that can help the investigator make an informed choice. These approaches have a shared goal: to minimize the threat of segmentation errors to measurement validity while maximizing retained sample size (when images are flagged for exclusion). A valid segmentation does not require perfection (i.e., without any error); some segmentation errors that are randomly distributed across slices and cases will introduce statistically negligible bias and thereby not affect validity of analyses with imperfect data. Therefore, choices for remediation of automated segmentation errors are about minimizing the greatest threats to validity and maintaining the efficiency (i.e., reduced time required) of automated methods.

#### Approaches to Re-segmentation of MR images and Recommendations

Commonly used methods for automated segmentation of the hippocampal subfields are often based on segmentation atlases that are validated against manual segmentations in a particular data set (e.g., Bender et al., 2018; Yushkevich et al., 2015b). Some software tools offer choices of segmentation atlases (e.g., ITK-SNAP, www.itksnap.org; Yushkevich et al., 2006). It is often difficult to predict the performance of these tools prior to applying them on a new data set. When selecting a segmentation atlas, an investigator should consider if the protocol has been validated in similar samples as the one to be processed (Wisse et al., 2020). Poor segmentation quality across the majority of a data set may indicate the specific segmentation atlas or protocol is not suitable. However, even when investigators select segmentation atlases that are appropriate to their studied population and ensure MR images are of sufficient quality, underperformance of the automated segmentation atlas can still occur and result in a high degree of error. This has been reported in the literature by investigators using validated segmentation atlases with MR images collected from both 3T and 7T magnets. For example, Wisse et al.’s (2017) segmentation atlas resulted in consistently smaller automatically segmented volumes of certain hippocampal subfields compared to manual segmentation. Another example is errors in the automated segmentation using the Bender et al. (2018) atlas that primarily occurred at the most anterior and posterior aspects of the hippocampal body. One solution to this problem is manual intervention. However, practical factors including rater expertise and time may be untenable, especially in large data sets. Thus, re-segmenting data with an alternative segmentation atlas or modified parameters is one approach investigators can take when severe errors are prevalent.

A decision to exclude all existing segmentations may be made if the investigator finds that a large number of scans fail to segment, a large number of severe segmentation errors affect the majority of the sample, or there is systematic bias in the segmentation errors with demographic features of the sample.

We recommend considering re-segmentation when segmentations fail on more than 40% of the data set, severe errors that threaten measurement validity are present in more than 40% of slices counted across all images, or when failed segmentations and major errors are systematically correlated with a variable of interest. This recommendation is based on guidelines on tolerance of data loss (see Little et al. (2014), McNeish (2017), and Raykov (2005) in-depth reviews, and section “Approaches to Data Exclusion and Recommendations” below for brief review of missing data tolerance and application to neuroimaging studies). If re-segmentation using a different segmentation atlas is not feasible or is unsuccessful, investigators may modify software-specific parameters and re-segment with the original segmentation atlas.

If a given segmentation software does not allow for parameter adjustment, then there may be other ways to correct for mis-segmentations. For example, in ASHS, frequent large mis-segmentations have been found to be caused by a subject’s T_1_-weighted scan not being properly co-registered to the T_1_ template or to their T_2_-weighted scan. This problem in registration is often due to the subject’s neck being too visible in their T1 scan, or due to the header information being incorrectly recorded in their T_1_- and T_2_-weighted scans, which can lead to problems in the initialization of the registration of the T_1_ and T_2_ scans. The solution to the first case is to trim the neck in the T_1_-weighted scan (which ASHS has a script for). In the second case, the T_1_- and T_2_- weighted scans can be run with a set of rigid registration parameters in ASHS. The other times ASHS fails are typically related to poor image quality (addressed in prior sections) or if the segmentation protocol is very distinct from the target data (i.e., there is a mismatch between the data used to develop the segmentation atlas and the data to be segmented).

Following the re-segmentation of hippocampal subfields, investigators should repeat the QC procedure on the re-segmented data. While not perfect, re-segmentation has been shown in practice to be a reasonable approach to retaining a greater proportion of data or reducing the number of errors that require further manual intervention. If issues of segmentation quality are not reduced following re-segmentation, investigators need to determine if manual correction or data exclusions are warranted. Final atlas selection and relevant parameters should be reported in the methods sections accompanying the data analysis. The trial-and-error of methods selection and atlas evaluation in the sample may be additionally helpful to the filed as we continue to refine our neuroimaging and segmentation procedures.

#### Approaches to Manual Correction of Automated Segmentation and Recommendations

With rater expertise and available time, errors in automated hippocampal subfield segmentation can be corrected manually. Manual intervention was reported by 43% of respondents as always being used to correct identified errors (the decision of *which* errors to correct differed between investigators), sometimes being used to correct identified errors by 31% of respondents (i.e., decisions on manual correction differed across data sets for the same investigator), and not being used to correct identified errors by 26% (i.e., errors were consistently uncorrected). Selective manual correction of only some errors, such as removing an obvious mislabel (e.g., cerebrospinal fluid) or adding labels to voxels that should have been labeled originally, is a viable approach to addressing severe segmentation errors while preserving benefits of automated segmentation efficiency, especially when working with large data sets. The severity ratings of segmentation errors made in prior QC steps are used to determine where manual intervention is applied (e.g., Canada et al., 2023). In this practice, manual corrections are made according to the segmentation protocol that was used to generate the segmentations which allows all images to be segmented using the same protocol regardless of whether it was corrected. Therefore, rater expertise in the segmentation protocol is a prerequisite of this approach.

###### Breakout Box 1: Considerations of bias and error.

Because human raters are prone to error, especially in the absence of clear rules, high reliability of a manual correction procedure must be established. Human error can be introduced to the measurement even with highly reliable correction procedures. The amount of human error introduced can be indexed by the ICC departure from 1.0 (because automated segmentation without human intervention shows high consistency) and should be both small and randomly distributed in order to mitigate bias. Unbiased error is supported if the frequency of corrected segmentations is not correlated with demographic features. If segmentation errors correlate with sample characteristics it leads to systematic bias in the measurement, even if the error is small. For example, in aged brains the loss of gray-white matter contrast is common, which could cause more segmentation errors for older brains as compared to younger ones for a given automated atlas.

Similar to our prior recommendations, human error should be minimized during the correction procedure by ensuring consistent decisions about data treatment and reliable error correction. However, as reflected by the responses of those surveyed, currently only 23% of respondents assess reliability of corrections to hippocampal subfields. As a best practice, we recommend establishing inter- or intra-rater reliability of the measurements following corrections in a subset of scans with errors before corrections are applied to the full data set. Specifically, we recommend reporting in the methods on good agreement in volume measures (with ICC > .85; Koo & Li, 2016) and spatial overlap (with DSC > .70; Zijdenbos et al., 1994) by making corrections on the same subjects to be compared between raters, or in the case of a single rater, with themselves after a delay. In addition, we recommend reporting the proportion of cases that were corrected or re-segmented using adjusted segmentation software parameters.

#### Approaches to Data Exclusion and Recommendations

Maintaining all cases in the collected sample is a top priority for external validity; however, there are instances where the segmentation errors cannot be remediated by re-segmentation or manual correction and the suitability of the MR images for hippocampal subfield segmentation may be deemed insufficient. In these instances, subfield segmentations from the original segmentation atlas are retained and the choice to exclude cases with errors that threaten measurement validity is reasonable. Among the investigators surveyed, 43% always excluded cases based on QC procedures, 31% sometimes excluded them, and 14% never excluded them despite segmentation errors. In most instances, cases with severe error are excluded from further analysis while cases with small or moderate errors are retained. Following from the QC decisions that vary by subregion and hemisphere, portions of the measured structure might be excluded while other parts are retained for the case (e.g., excluding left hemisphere but retaining the right hemisphere; or excluding hippocampal head measurements while retaining measurements in hippocampal body).

When cases are excluded, the criterion for exclusion and number of cases excluded should be reported and noted as a limitation. Exclusion of cases contributes to overall data loss and impacts statistical analysis and interpretation. Excluded cases are missing data; therefore, the same statistical considerations for randomness should apply. The loss of data due to exclusion may meaningfully reduce statistical power for subsequent hypothesis testing. The statistical literature offers a good overview of missing data considerations (Little et al., 2014; McNeish, 2017; Raykov, 2005), which we will review briefly for applied neuroimaging studies.

The decision about parameter estimation under the condition of missing data is informed by three criteria: the planned statistical estimation method, the amount of missing data, and the randomness of missingness. We have emphasized detection of errors during QC to maximize validity of the hippocampal subfield measurements, but we also must consider if the retained sample upon completion of QC still represents the study population. Many of the common analyses in applied neuroimaging studies are based on cases with complete data. Listwise or pairwise deletion of cases of missing data (in this case by exclusion) will provide unbiased estimates of the population-level effect only when data are missing completely at random (Baraldi & Enders, 2010). There are alternative approaches including multiple imputation or latent modeling with incomplete data (see Little et al. (2014) for an overview) that can maximize external validity and have additional benefits to statistical power, but these also rely on data missing at random to provide unbiased estimates. To support the conclusion of data missing at random, formal test statistics (e.g., Little’s chi-square test) can indicate for randomness and the frequency of data loss due to QC should be negligibly correlated with demographic or study variables of interest, as shown with descriptive statistics or logistic regression. In practice, however, the assumption of data missing completely at random in QC is difficult to meet as more severe cases are more likely to fail QC, which underscores the importance of re-segmentation and manual correction options to retain as much data as possible. Based on current recommended practices for statistical tolerance of missing data (Little et al., 2014; McNeish, 2017; Raykov, 2005), we recommend that no more than 40% of data should be lost to QC issues collectively.

## Data Screening

### Description of the problem

Following QC of hippocampal subfield segmentation labels, an important additional step in the process is statistical data screening prior to hypothesis testing. Data screening helps provide assurance that the data included for analysis is accurate, meets assumptions of the planned statistical analyses and promotes rigorous best practices.

### Review of Current Approaches and Recommendations

Data screening following other steps in the QC procedure is an important last check point to identify errors such as statistical outliers in the data and is an approach taken by 33% of the respondents. Although prior QC steps discussed will facilitate the identification of the majority of errors, some errors may be overlooked, or investigators may choose to conduct a subset of the reviewed procedures. Thus, data screening of measurement values provides a secondary check of the QC procedures. A common first step in data screening is the inspection of univariate descriptive statistics. This includes examining data for out-of-range values (e.g., implausibly small or large volumes), inspecting the means and standard deviation for plausibility, and assessing the presence of univariate outliers. This step was included in the QC steps of most respondents, with 54% examining segmentation values for outliers within the sample. Screening procedures for out-of-range values and outliers are often repeated for volumes adjusted for intracranial volume that are intended for planned analysis, which is a sample-specific procedure that can modify the relative rank of a case relative to its sample distribution.

As an initial data screening approach, an investigator can examine correlations of a regional measure between hemispheres: for many populations, high consistency is expected and cases that deviate from the diagonal of the scatterplot would be candidates for review. Outlier detection is the highest priority in data screening related to QC best practices. With a reasonable approximation of a normal distribution, univariate outliers can be determined by z-scores exceeding |3.29|. However, the majority of statistical methods used in applied neuroimaging studies are more sensitive to multivariate outliers than univariate deviations. For hippocampal subfield measurements, assessing multivariate normality by quantile-quantile (Q-Q) or probability-probability (P-P) plots is recommended. Multivariate outliers can be detected by Mahalnobis distance with a cut-off of a critical chi-square (degrees of freedom = number of variables; alpha = 0.001), in addition to assessments of outliers in regression residuals during hypothesis testing (Tabachnick & Fidell, 2019). Decisions to remove outliers from analyses are another instance of missing data, which contributes to the overall consideration of tolerance in the statistical design discussed above. It is important to note that investigators should be aware of individual differences in data sets (e.g., cases of severe neurodegeneration) when assessing outliers. Here we have described the use of statistical outlier detection as a means to identify cases for QC review, but extreme values are realistically plausible especially in the scenario of a severe disease stage. After review of an outlier case, the investigator may determine it meets all criteria that we have reviewed to retain in the sample—it is, in fact, an accurate measurement— and proceed with analysis following standard practices with outlier values.

## Special considerations for longitudinal data sets

Different procedures can be used to ensure continuity in longitudinal studies. One approach is segmentation of data collected at each measurement independent of prior timepoints. During QC, raters can refer to all available timepoints when evaluating possible segmentation errors. This type of approach has produced high test-retest consistency and supports measurement invariance over longitudinal assessments (Homayouni et al., 2021). In addition, when applied to manually segmented data, raters can be blinded to timepoint to ensure no bias is introduced.

An alternative approach is to study longitudinal change in subfields using deformation-based morphometry (Das et al., 2012). In this approach, the change in volume is combined by performing registration between MRI scans at different timepoints, and only one timepoint needs to be segmented to obtain subfield-specific measures of change. While this approach reduces the burden of QC of segmentations, it does require QC of the MRI scans at each timepoint separately and of the pairwise registrations. However, QC of pairwise registrations can be done using a semi-automatic method by examining registration quality metrics (e.g., intensity cross-correlation in the MTL region after registration) and focusing manual QC procedures on image pairs where that metric is more than 2.5 standard deviations from the mean (Xie et al., 2020).

It is important to note that although there are differences in the approach for longitudinal segmentation, the QC procedures we have reviewed above apply similarly to data at all timepoints (Shaw et al., 2020). When datasets are large, the general quality of segmentation and potential extent of segmentation errors can be approximated from the procedures applied to a subset of cases that are selected at random. This approach benefits from the same principle of sampling in statistical analysis: if the subset is randomly selected and representative of the sample features, then the frequency and severity of segmentation errors is expected to generalize to the whole sample. Because the goal of QC is to minimize systematic measurement error, and the goal is not perfect data *per se*, this subset evaluation may identify that major threats to validity are infrequent and therefore the investigator retains the data for further analysis. In the event that severe image artifacts and severe segmentation errors are identified with high frequency in a subset, the investigator can weigh the further time investment to screen the dataset in more detail.

## Reporting QC Procedures

All data screening procedures and decisions for data conditioning or exclusion should be described in publications. Describing the amount of missing data and reasons for data loss (i.e., poor image quality, segmentation failure, segmentation error, or statistical outlier) is paramount for determining the external validity of the analysis based on the representativeness of the retained sample.

### Guidelines for implementing and reporting QC methods*

Note (*) that we review options for QC procedures and provide guidelines for the reporting and implementing of QC that the investigator can use to inform their choice of QC steps. The recurring recommendation for best practice across all possible QC steps is reporting in publications which steps were implemented and with enough detail so that readers can sufficiently understand the decisions made and amount of data affected.

| QC Step | Recommendations | What to report: |
| --- | --- | --- |
| Assessment of MR Image Quality | • Rating scale for image quality and landmark visualization<br>• Optional: SNR/CNR | • Criteria for exclusion (e.g., rating scale, yes/no)<br>• Number of raters<br>• Rater reliability (inter and/or intra) of scans (kappa statistic)<br>• Number (percentage) of scans excluded<br>• If exclusion correlated with variables of interest |
| Assessment of Segmentation Quality | • Rating scale of severity of error<br>• Determine amount of data loss due to segmentation failure (less than 40 recommended%) | • If segmentations were reviewed and criteria for judgment<br>• Number of raters<br>• Rater reliability (inter and/or intra) of manual segmentations (kappa statistic)<br>• Number (percentage) of segmentations failed<br>• Potential covariates of segmentation error frequency or exclusion |
| Options for Error Remediation |  |  |
| Manual Correction of Automated Segmentation | • Establishing reliability of correction procedure<br>• Correction of only severe errors to reduce human bias | • If segmentations were corrected, what procedure was used<br>• Number of raters<br>• Rater reliability (inter and/or intra) of the correction procedure for measurement of interest: volume (ICC2) or mask (DSC coefficient)<br>• Number (percentage) of segmentations corrected |
| Data Exclusion | • Define specific parameters for exclusion<br>• Assessment of data loss due to exclusion (percent)<br>• Assessment of missing data assumptions | • Number (percentage) of segmentations excluded<br>• Chi-Square test for MCAR<br>• Potential covariates of exclusion |
| Data Screening | • Assess distribution of data<br>• Examine and exclude outliers<br>• Report criterion used and data excluded | • Univariate outlier detection<br>• Assessment of missing data<br>• Multivariate outlier detection (e.g., Mahalanobis distance)<br>• High correlation between hemispheres<br>• Longitudinal consistency within hemisphere (ICC3, DSC) |

## Conclusion

In the ever-growing field of hippocampal subfields neuroimaging research, ensuring accurate measurement is vital to progress our understanding of brain structural and functional correlates, lifespan development, and neuropathology. QC is an essential part of ensuring valid results by promoting accurate hippocampal subfield segmentation and retaining all eligible data for analysis. In addition, recent advancements in neuroimaging have given investigators the unique opportunity to study hippocampal subfields with greater precision than ever before in pursuit of the ultimate goal of drawing robust scientific conclusions that meet a high standard of quality.

Using the literature and our findings from a survey of investigators with experience segmenting hippocampal subfields, we summarized the threats to segmentation accuracy, reviewed common methods for QC, and made recommendations for best practices and reporting of QC in publication. In the following section we highlight broader impacts of ensuring quality data on neuroimaging and clinical research.

The importance of QC is underscored by goals related to investigating and validating structure-function relationships. The study of hippocampal subfields is a unique *in vivo* endeavor as a major focus is the ability to understand the nuanced roles of hippocampal function in human cognition and forward translation from animal models of development, aging, and disease. For example, determining the impact of normative and pathological aging on hippocampal structure and related memory abilities requires reliable measurements of both cognition and brain that allow accurate conclusions from hypothesis testing. Moreover, validity is critical in supporting the clinical translation of hippocampal subfields measurement to biomarkers of healthy development, neurodegenerative disease, and neurodevelopmental disorders. The requisite of interpreting hippocampal subfields as biomarkers is confidence that the measures used reflect the underlying tissue structure and its changes over the course of time.

Further, QC of hippocampal subfield segmentations is essential to promoting reproducibility of findings and methods because the subfields are small regions that share boundaries, increasing the risk of errors with serious consequences to validity. While we focused on hippocampal subfield segmentation in this manuscript to highlight the importance of QC, it should be noted that it is within the larger context of measurement reliability and validity in applied neuroimaging. Thus, the QC practices we recommend here play a two-fold role in supporting reproducible neuroimaging research broadly. First, using best practices in QC supports reproducibility in neuroimaging research by focusing on transparent reporting of the methods and decisions made. Reporting decisions made in the treatment of data from post-acquisition to analysis contextualizes the results of a given study within the larger literature and provides a path forward for others to implement similar approaches in the study of hippocampal subfields. Second, harmonization of methods across laboratories, including QC procedures, is essential to reproducible research. The Hippocampal Subfields Group (HSG) is leading harmonization efforts, including a harmonized set of definitions for subfield segmentation (Olsen et al., 2019; Wisse et al., 2017; Yushkevich et al., 2015a). These efforts have highlighted another gap in the available literature, best practices for QC. We attempt to address this gap by providing recommendations that will apply to any segmentation approach, including a harmonized procedure that is in development. The need for QC remains critical, especially with harmonization efforts, as site- and sample-specific errors that affect image quality and segmentation accuracy can occur even with a harmonized segmentation protocol. With possible errors occurring for the many reasons discussed in this paper, the application of QC procedures aids in the harmonization of data, ensuring transparent and appropriate decision making across sites. With repeated calls for large, representative data sets in the neuroimaging field, transparent knowledge and reporting of decisions made when segmenting, correcting, and ultimately retaining measures of hippocampal subfields are facilitated by the QC procedures recommended here. In addition, it is critical for research groups to share their data and QC procedures in their publications in order to aid in the further development of robust automatic QC tools (e.g., artificial intelligence-based QC).

Retaining all eligible data is a priority for external validity to address substantial limitations in sample representation and inferences on development, aging, and neuropathology. As we have discussed, there is the potential for bias in the occurrence of errors. For example, the risk of threats to validity is increased when applying automated methods with segmentation atlases developed in healthy populations to clinical populations or vice versa. Identifying and correcting errors in hippocampal subfield segmentations ensures measurement validity is acceptable across different subpopulations. Thus, implementing QC to identify and correct data errors increases not only the validity of the measures but also the validity of the inferences to the larger population and subsequently the implications for clinical practice and public health.

## Acknowledgements

We thank the respondents of the Hippocampal Subfields Group Best Practices Survey for their invaluable insights including Étienne Aumont, Pedro Renato de Paula Brandão, Nicholas Christopher-Hayes, Jordan DeKraker, José Carlos Delgado-González, Maria-Eleni Dounavi, Matt Dunne, Nicole Gervais, Juan Domingo Gispert, Maria del Mar Arroyo Jimenez, Anne Maass, Elizabeth McDevitt, Jennifer Park, María Pilar Marcos Rabal, Tracy Riggins, Carlos de la Rosa-Prieto, Monica Sean, Christine Smith, Alina Tu, and Davis Woodworth. We also thank Carl Hodgetts and Kim Nguyen for providing landmark visibility quality ratings.

