## Supplementary Material for "A (Sub)field Guide to Quality Control in Hippocampal Subfield Segmentation on High-resolution T_2_-weighted MRI"

^+^ denotes shared first-author position.

^#^ denotes shared senior-author position.

*Corresponding Authors:

Kelsey L. Canada, Institute of Gerontology, Wayne State University, Detroit, MI, USA; Ana M. Daugherty, Institute of Gerontology and Department of Psychology, Wayne State University, Detroit, MI, USA,

Affiliations

^1^ Institute of Gerontology, Wayne State University, Detroit, MI 48202

^2^ Department of Psychology, University of Toronto, Toronto, Ontario, Canada

^3^ Rotman Research Institute, Baycrest Health Sciences, Toronto, Ontario, Canada

^4^ Department of Clinical Sciences Lund, Lund University, Lund, Sweden

^5^ Department of Neurobiology and Behavior, University of California, Irvine, CA 92697

^6^ Department of Psychiatry and Behavioral Sciences, Johns Hopkins University School of Medicine, Baltimore, MD 21287

^7^ German Center for Neurodegenerative Diseases (DZNE), Magdeburg, Germany

^8^ Dementia Research Centre, Department of Neurodegenerative Disease, UCL Queen Square Institute of Neurology, University College London, London, UK

^9^ Centre for Cognitive and Clinical Neuroscience, Division of Psychology, Department of Life Sciences, College of Health, Medicine and Life Sciences, Brunel University London, London, UK

^10^ Department of Psychology, San Jose State University, San Jose, CA 95192

^11^ School of Psychology, University of Sydney, Sydney, Australia

^12^ INSERM UMR-S U1237, Physiopathology and Imaging of Neurological Disorders (PhIND), Institut Blood and Brain @ Caen-Normandie, Caen-Normandie University, GIP Cyceron, France

^13^ Brain Imaging Centre, Research Centre for Natural Sciences, Eötvös Loránd Research Network (ELKH), Budapest, Hungary

^14^ Institute of Psychology, ELTE Eötvös Loránd University, Budapest, Hungary

^15^ Center for Lifespan Psychology, Max Planck Institute for Human Development, Berlin, Germany

^16^ Memory and Aging Center, Department of Neurology, Weill Institute for Neurosciences, University of California, San Francisco, CA 94158

^17^ Department of Radiology, University of California, San Francisco, CA 94143

^18^ Center for Imaging of Neurodegenerative Diseases, San Francisco VA Medical Center, San Francisco, California 94121

^19^ Department of Psychology, Stony Brook University, Stony Brook, NY 11794

^20^ Department of Bioengineering, University of Pittsburgh, Pittsburgh, PA 15213

^21^ School of Electrical Engineering and Computer Science, The University of Queensland, Brisbane, Australia

^22^ Department of Psychology, Stanford University, Stanford, CA 94305

^23^ Department of Psychiatry and Behavioral Health, The Ohio State University Wexner Medical Center, Columbus, OH 43210

^24^ Clinical Memory Research Unit, Department of Clinical Sciences, Malmö, Lund University, Sweden

^25^ Penn Image, Computing and Science Laboratory, Department of Radiology, University of Pennsylvania, Philadelphia, PA 19104

^26^ Department of Psychology, Wayne State University, Detroit, MI 48202

^27^ Michigan Alzheimer’s Disease Research Center, Ann Arbor, MI 48105

*To reference this document, cite: XXX*

**Survey Detailed Response Information**

**Section 1: Information about respondents and MRI methods**

| **Table S1.1** Demographics (N = 37) | | | |
| --- | --- | --- | --- |
| **Country** | Count (proportion) | Country | Count (proportion) |
| **USA** | 17 (45.9 %) | **Great Britain** | 2 (5.4 %) |
| **Canada** | 5 (13.5 %) | **Brazil** | 1 (2.7 %) |
| **Spain** | 5 (13.5 %) | **New Zealand** | 1 (2.7 %) |
| **Germany** | 2 (5.4 %) | **Hungary** | 1 (2.7 %) |
| **Sweden** | 2 (5.4 %) | **France** | 1 (2.7 %) |

| Table S1.2 Populations studied across research groups (N = 37) | | |
| --- | --- | --- |
| Population | | Count (proportion) |
| Healthy | | 33 (89.2 %) |
| Cognitive Impairment (SCI/MCI) | | 21 (56.8 %) |
| Alzheimer’s disease | | 19 (51.4 %) |
| Epilepsy | | 5 (13.5 %) |
| White matter hyperintensities | | 4 (10.8 %) |
| Hypoxia/anoxia | | 3 (8.1 %) |
| Depression | | 2 (5.4 %) |
|  | Other *(Respondent provided answer)* |  |
| Parkinson’s disease | | 2 (5.4 %) |
| Development (incl. developmental disorders) | | 2 (5.4 %) |
| Traumatic Brain Injury | | 2 (5.4 %) |
| PTSD | | 1 (2.7 %) |
| Frontotemporal dementia | | 1 (2.7 %) |
| Psychosis. Traumatic Brain, Parkinson’s Disease | | 1 (2.7 %) |
| Chronic pain population | | 1 (2.7 %) |
| Motor Neuron Disease (ALS) | | 1 (2.7 %) |
| Preclinical Alzheimer’s disease | | 1 (2.7 %) |

| Table S1.3 Age groups studied across research groups (N = 37)* | |
| --- | --- |
| Age group | Count (proportion) |
| Children (0-10) | 6 (16.2 %) |
| Adolescents (10-18) | 4 (10.8 %) |
| Young adults (18-30) | 19 (51.4 %) |
| Middle-aged adults (35-55) | 21 (56.8 %) |
| “Young” older adults (55-75) | 32 (86.5 %) |
| “Old” older adults (>75) | 30 (81.1 %) |
| **answers not mutually exclusive* |  |

| **Table S1.4** Magnetic field strength used across research groups (N = 37) | |
| --- | --- |
| **Magnetic field strength** | Count (proportion) |
| **1.5T** | 5 (13.5 %) |
| **3T** | 35 (94.6 %) |
| **4T** | 2 (5.4 %) |
| **7T** | 12 (32.4 %) |

| Table S1.5 MRI Sequence used across research groups (N = 37) | | |
| --- | --- | --- |
| MRI Sequence | | Count (proportion) |
| T1-weighted images (1mm3) | | 21 (56.8 %) |
| T2-weighted images | | 28 (75.7 %) |
| Fluid Attenuated Inversion Recovery (Flair) | | 1 (2.7 %) |
| Proton Density (PD) | | 4 (10.8 %) |
|  | Other *(Respondent provided answer)* |  |
| T1-weighted images (0.5-0.8mm3) | | 1 (2.7 %) |
| Inversion Recovery high resolution (0.4x0.4x2, 0mm) | | 1 (2.7 %) |

**Section 2 – General use and QC of MR images**

| **Table S2.1** How important are quality control procedures? (N = 37) | |
| --- | --- |
| **Response** | Count (proportion) |
| **Very important** | 28 (75.7 %) |
| **Important** | 2 (5.4 %) |
| **Fairly important** | 7 (18.9 %) |
| **Slightly important** | 0 (0 %) |
| **Not important at all** | 0 (0 %) |

**Note*: Very important and important collapsed for reporting in manuscript.

| Table S2.2 Do you review the quality of MR images? (N = 37) | |
| --- | --- |
| Response | Count (proportion) |
| Yes | 33 (89.2 %) |
| No | 4 (10.8 %) |

| Table S2.3 During QC of MR images, which methods do you use? (N = 33) | |
| --- | --- |
| Response | Count (proportion) |
| Visual inspection | 33 (100 %) |
| Outlier detection | 11 (33.3 %) |
| Automated QC procedures | 5 (15.2 %) |
| Visualization tools | 6 (18.2 %) |

| Table S2.4 During QC, do you exclude images based on quality? (N = 33) | |
| --- | --- |
| Response | Count (proportion) |
| Yes | 31 (93.9 %) |
| No | 2 (6.1 %) |

| Table S2.5 Describe exclusion criteria or (if no) motivate retaining all images (N = 26) | | |
| --- | --- | --- |
| Problems discussed | Count | Comments |
| Artifacts |  |  |
| Motion artifacts | 13 | Visualized as “rings” (1), only when multiple images are available (1), “study specific cut-off” (1) |
| Ringing artifacts | 3 |  |
| Banding artifacts | 1 |  |
| Ghosting artifacts | 1 |  |
| Blurring artifacts | 3 | Often caused by motion artifacts, but other blurring mechanisms might be referred to as well such as heart movements. |
| Visibility |  |  |
| Border visibility problems | 11 | Dark band/SRLM, gray matter//white matter borders. |
| Coverage problems | 2 | E.g., Hippocampus “cut off” in images. |
| Axis deviation | 1 | Excluding when the deviation from the axis along the hippocampal body is too large. |
| Quantitative |  |  |
| Measurement outliers | 2 | Second stage quality check prior to analysis. One indicated using “volume averages” for this, the other did not specify. |
| SNR and/or CNR | 2 |  |
| Hemispheric agreement | 1 |  |
| Rating systems | 2 | (a) Subjective general assessment categorizing as “Viable for semi-automatic segmentation”, “Viable for manual segmentation” or “Not viable at all”.  (b) “Usable”, “Questionable”, “Not usable” determined using qualitative judgment and SNR. |
| Other information |  |  |
| Subjective judgment | 4 | E.g., assessing distortion, excluding if manual segmentation is unfeasible, see rating systems (a). |
| Analysis-dependent | 1 | (a) Whole brain vs. regional vs. voxel-based, (b) group-comparison vs. single cases. |

**Note*: Motion categories collapsed in manuscript.

**Section 3 – QC of Hippocampal Subfield Segmentations**

| Table S3.1 Do you review the quality of hippocampal subfield segmentations regardless of a manual or automated approach? (N = 37) | |
| --- | --- |
| Response | Count (proportion) |
| Yes | 35 (94.6 %) |
| No | 2 (5.4 %) |

| Table S3.2 What approach(es) do you take for the segmentation of hippocampal subfields?  (N = 35) | |
| --- | --- |
| Response | Count (proportion) |
| Automated segmentation | 29 (82.9 %) |
| Manual segmentation | 17 (48.6 %) |
| Automated supplemented with manual edits | 1 (2.9 %) |
| Template based segmentation | 1 (2.9 %) |

**Note*: Template based segmentation is considered automated segmentation and reported as such in manuscript.

| Table S3.3 Technical specifications and protocols/atlases | |
| --- | --- |
| Software | Count |
| ASHS | 23 |
| *ASHS 1.0.0* | *2* |
| *ASHS 2.0.0* | *4* |
| *ASHS (unspecified version)* | *17* |
| ITK-SNAP | 17 |
| *ITK-SNAP 3.6* | *1* |
| *ITK-SNAP 3.8* | *5* |
| *ITK-SNAP (unspecified version)* | *11* |
| Freesurfer | 14 |
| *Freesurfer 6* | *3* |
| *Freesurfer 7(.1/.2)* | *4* |
| *Freesurfer (unspecified version)* | *7* |
| VolBrain | 2 |
| FSL | **2** |
| *FSLView 4.0* | *1* |
| *FSLeyes 6.0.0-6.0.4* | *1* |
| HippUnfold v1.0.0-1.2.0 | **1** |
| MAGetbrain | **3** |
| LASHiS | **1** |

| Table S3.3.1 Technology used for segmentation. For manual segmentation, please note the type of tablet used OR indicate if tracing is done on computer with a mouse. For automated segmentation please answer "auto." (N = 35) | | |
| --- | --- | --- |
| Technology | Count | Comments |
| Automated | 21 | Counted as those who mention auto, note that many also report doing manual segmentations or manual correction of automatic segmentations. |
| Computer with mouse | 11 |  |
| Tablet with stylus | 4 |  |
| Computer with trackpad | 1 |  |
| Tablet with mouse | 1 |  |
| ITKSnap | 1 | For completion. |

| **Table S3.4** Do you correct segmentations (i.e., manually edit) based on your quality control procedures? (N = 35) | |
| --- | --- |
| **Response** | Count (proportion) |
| **Yes** | 15 (42.9 %) |
| **No** | 9 (25.7 %) |
| **Sometimes** | 11 (31.4 %) |

| Table S3.5 Please provide a brief description of the procedures you use to make corrections to your segmentations. (N = 22) | | | |
| --- | --- | --- | --- |
| Segmentation errors mentioned | | Count | Comments |
| Oversegmentation errors | |  |  |
|  | Lateral oversegmentation (e.g., choroid plexus included as CA1) | 2 |  |
|  | Voxels extending into fimbria | 1 |  |
|  | Voxels scattered around subfields | 1 |  |
|  | Random blobs not attached to the hippocampus | 1 |  |
|  | Incorrect inclusion of ventricles | 1 |  |
|  | Segmentations “jumping” over to brainstem | 1 |  |
|  | Oversegmentation of the MTL cortices | 1 |  |
| Undersegmentation errors | |  |  |
|  | Lateral undersegmentation of the CA1 | 1 |  |
|  | CA1 + CA2 + Subiculum | 1 | Only if there are several slices in a row |

| Table S3.5 continued. | | |
| --- | --- | --- |
| Overlap between subfield | 1 |  |
| Mislabeling | 1 | Preliminarily identified by trained lab member, corrected by more experienced member. |
| Discontinuous labeling | 1 |  |
| Unusual shapes | 1 |  |

| **Table S3.6** Please provide a brief description of how you evaluate the quality of your segmentation correction. (N = 19) | |
| --- | --- |
| **Response** | **Comments** |
| **Visual inspection** | - Ensuring only gray matter is included in labels - Ensuring dark band separates DG and SUB/CA - Clear separation of digitations/dentations |
| **Using several raters or some form of group revision by members of the lab** | - Some form of intra/inter-rater analysis to check for quality of correction (e.g., DICE or “Inter-rater ICC on hippocampal volume”). - Some also report using reliability estimates acquired on training data to estimate the degree of “human error” and as a quality check in analysis. |
| *Other comments* |  |
|  | - Applying minor corrections, leaving the evaluation of quality up to the editor of the segmentation. - Comparing the corrected to the original segmentation - Failed segmentations can be identified and resolved. As a result, the scan is re-submitted for automated segmentation |

| Table S3.7 Do you exclude segmentations based on your quality control procedures? (N = 35) | |
| --- | --- |
| Response | Count (proportion) |
| Yes | 19 (54.3 %) |
| No | 5 (14.3 %) |
| Sometimes | 11 (31.4 %) |

**Table S3.8** Please provide a brief description of the quality control procedures you use for the exclusion of your segmentations. (N = 5)

**Comments**

Many visually inspect results, looking for larger errors or obvious mistakes.

Generally, larger errors mean a full subject exclusion.

Inability to confidently appraise or correct segmentations (due to image quality).

Outliers based on volume SDs or the extent of error (x mm) along the longitudinal axis of the HC.

Using protocols or manuals to make QC decisions. These include 1) MAGeT-related QC (https://github.com/CoBrALab/documentation/wiki/MAGeT-Brain-Quality-Control-(QC)-Guide), 2) an automated QC that’s part of HippUnfold (DICE overlap with a coarser method (deformable registration), 3) MRIQC (Esteban et al., 2017).

Deploy a rating system that will help make the exclusion decision.

| Table S3.9 Do you make different decisions about your exclusions or corrections depending on:  (N = 35) | |
| --- | --- |
| Response | Count  (proportion) |
| Sample size | 9 (25.7.3 %) |
| Uniqueness of population | 11 (31.4 %) |
| Bias in segmentation errors | 11 (31.4 %) |
| I do not make different decisions depending on these factors. | 17 (48.6%) |
| *Other:* Yes. balancing how much information the segmentation adds vs. noise | 1 (2.9%) |
| *Other:* At least consciously not | 1 (2.9%) |

| Table S3.10 Which of the following do you use during the quality control of your segmentations? (N = 35) | |
| --- | --- |
| Response | Count (proportion) |
| Inter/Intra-rater reliability of quality ratings | 11 (31.4 %) |
| Inter/Intra-rater reliability of manual segmentations (e.g., during protocol establishment and/or training of new raters) | 13 (37.1 %) |
| Inter/Intra-rater reliability of segmentation corrections | 8 (22.9 %) |
| Cross-validation of automated segmentations to manual segmentations of hippocampal subfields | 11 (31.4 %) |
| Outlier detection of segmentation values | 19 (54.3 %) |
| None of the above nor "Other" | 7 (20 %) |
| *Other:* “I will need to but have not yet done so” | 1 (2.9 %) |
| *Other:* “Visual inspection vs. anatomy” | 1 (2.9 %) |
| *Other:* “Visual inspection by an expert” | 1 (2.9 %) |

| Table S3.11 Do you use a scoring/rating system during the QC of segmentations? (N = 35) | |
| --- | --- |
| Response | Count (proportion) |
| Yes | 16 (45.7 %) |
| No | 15 (42.9 %) |
| Sometimes | 4 (11.4 %) |

| Table S3.12 Scoring system details | | |
| --- | --- | --- |
| Rating system | Count | Comments (Number of respondents) |
| Usable/Unusable | 4 | - Ensuring only gray matter is included in labels - Ensuring dark band separates DG and SUB/CA - Moved away from using a 5-point scale because the information was not used (1) |
| Four-point scale | 3 | - Best – Good – Questionable – Reject (1) - Poor – Fair – Good – Great (1) - Include – 3D-check needed – Exclude – Exclude specific subfield - Unusable – Bad – Okay – Good (bad and unusable typically excluded) (1) |
| Three-point scale | 6 | - Passed – Borderline – Failed (1) - Only automated segmentations are scored for quality. Rated for severity on a 3 pt scale (1-least severe, 3-most severe) that is based on an estimated percentage of slice area that is affected by the error. (1) - Inclusion – Inclusion but with limitations – Exclusion. We exclude these cases when running sensitivity analyses (1) - Good – Poor – Unusable. "Poor" entails small errors that will likely not result in exclusion except for the most selective of analyses. (1) - Unusable – Problematic but usable – Usable (1) - Pass – Warning – Fail (1) |
| Five-point scale | 2 |  |
| Six-point scale | 1 |  |

| **Table S3.12.1** Is your scoring system slice-based or voxel-based (e.g., X number of pixels results in an Unusable rating, versus X number of slices have errors present)? (N = 15) | | |
| --- | --- | --- |
| **Response** | Count | Comments |
| **Slice-based** | 5 |  |
| **Voxel-based** | 2 |  |
| **Both** | 3 | ” …the presence of large (pixel) based errors in tandem with the consistency of errors (slices)” (1) |
| **Other** | 2 | ”Voxel-based for 3D, slice-based for 2D but still with some sense of volumetric measures. Of course, it depends on eventual analysis, so slice-based if particular slices are more important than total volume.” (1)  “It is based on approximated area, similar to number of pixels.” (1) |
| **Unsure** | 1 |  |
| **No scoring system** | 2 |  |

| **Table S3.12.2** Are segmentations reviewed by multiple raters? (N = 20) | |
| --- | --- |
| **Response** | Count (proportion) |
| **Yes** | 2 (10 %) |
| **No** | 6 (30 %) |
| **Sometimes** | 12 (60 %) |

| Table S3.13 Do you work with longitudinal data? (N = 35) | |
| --- | --- |
| Response | Count (proportion) |
| Yes | 19 (54.3 %) |
| No | 16 (45.7 %) |

| Table S3.14 How do you typically QC longitudinal data (e.g., do you QC data at each time point)? (N = 16) | | |
| --- | --- | --- |
| Response | Count | Comments |
| Each time point | 10 | - “Independently and blindly” (1) |
| *Special comments* | 1 | - “Only the baseline data is segmented and labels for follow up time points are created through registration warps. The idea is that the error introduced by registration are smaller than the error by segmentation. For longitudinal data the registration pairs are checked. In case of a large number of registration pairs (e.g. several 100s), metrics are used to select pairs with a likely bad registration + a certain percentage of random pairs. Pairs with bad registration are then excluded.” - “We only check 1 time point first (subject average). If an error is detected, all time points are checked and edited” - “We are still in the data acquisition phase of the longitudinal study” - “Raters are blinded to timepoint by double coding the MRI scan IDs. All decisions about scan quality, ranging and segmentation are made considering all time points. Ranging and manual segmentations are made while consulting all timepoints, and segmentations are completed together in one sitting. Automated segmentations may not be reviewed simultaneously for all timepoints, but decisions for exclusion and evaluation of bias are completed with time-nested data comparisons. Exclusion of cases based on outlier detection are made on a scan-by-scan basis, but implications for analysis as listwise or pairwise deletion are considered based on the planned longitudinal analysis. Data exclusion or statistical adjustment for bias/reliability are made with consideration of measurement invariance, sample representation and options for time-varying covariate adjustments.” - “Single subject template and after segmentation” |

| Table S3.15 Do you apply the procedures you’ve described to other anatomical segmentations? (N = 35) | |
| --- | --- |
| Response | Count (proportion) |
| MTL cortex | 19 (54.3) |
| Total hippocampus | 22 (62.9 %) |
| Hippocampal subregions (e.g., head, body, tail) | 24 (68.6 %) |
| I do not apply these procedures to other anatomical segmentations | 5 (14.3 %) |
| *Other: “ROIs”* | *1 (2.9 %)* |

| Table S3.16 Have you reported quality control procedures in prior publications? | |
| --- | --- |
| Response | Count (proportion) |
| Yes | 17 (45.9 %) |
| No | 20 (54.1 %) |

**Table S3.17** What resources/tools would need to be more readily available/accessible in order for you to be able to QC your data more/incorporate QCing more into your segmentation practices? (N = 25)

**Comments**

Protocol providing guidance: “Pictures/videos of step-by-step segmentation examples, including various types of hippocampal shape”

Automated methods: “Automatic MRI QC tools like MRIQC but for automatic segmentation.”

Better visualization tools: “Comparison with shape templates to detect outlier behavior could be useful for screening purposes, to optimize the process of manually inspecting the images.”

Happy as is: “I'm happy with ITK-Snap for QC checks and correction. If there was a reliable automatic tool, I would be happy to use it.”

**Table S3.18** Please note if there are specific problems you encounter when trying to QC your data.

**Comments**

Different Collateral Sulcus types (“though I know that it's not relevant for hippocampal subfields”)

Lack of statistical assessment in literature

Intra-individual variability (“…is a key factor which makes QC-ing less clear in some cases”)

Under-segmentation most common in automated segmentation

“Switching between PDFs of snapshots and ITK Snap can be tedious”

“Even among neurologists/radiologists hippocampal subfield anatomy is not as well-known as in the cortex”

Time consuming

Table S3.19 Are there other topics not covered in this survey you think are important to consider in the conversation about quality control to promote valid and reliable data? (N = 14)

Comments

“I think the biggest way to promote validity and reliability is a singular segmentation protocol, which I'm so grateful you all have been working on. I struggled most with how to segment subfields that look different across participants (i.e. varying # of digitations in the hippocampal head), so detailing "exceptions" to the protocol will be really helpful (if that's possible).”

“There is an interesting topic about this subject in a recent article in Hum Brain Mapp, from the ENIGMA consortium (https://www.ncbi.nlm.nih.gov/pmc/articles/PMC8805696/”)

“Harmonization across labs and populations”

“After getting my segmentations, I use them to mask my epi data, and then run quality control procedures (e.g., looking at mean voxel intensity and tsnr in each voxel in each subfield) to determine if some areas have more signal dropout than others, as these are areas that can be prone to dropout.”

“For automatic segmentation (e.g., ASHS), whether some editing of the segmentations during QC is performed - can be beneficial (in severe cases of mislabeling) or could add potential bias if not performed on all cases/datasets and introduce variability if multiple people are editing the automatic segmentations with different criteria”

“It may be best to ensure mesoscopic (ie. digitations/dentations) segmentation is correct before examining microscopic (ie. neuronal distributions) features”

“Quality control of the images per se. Are they of sufficient quality to be used for subfield segmentation?”

“Reliability vs accuracy, multiple acquisitions in a single session, better sequence/protocol choices at 3T, differences in 7T vs 3T, data quality for patient groups.”

**Additional Examples of MRI QC**

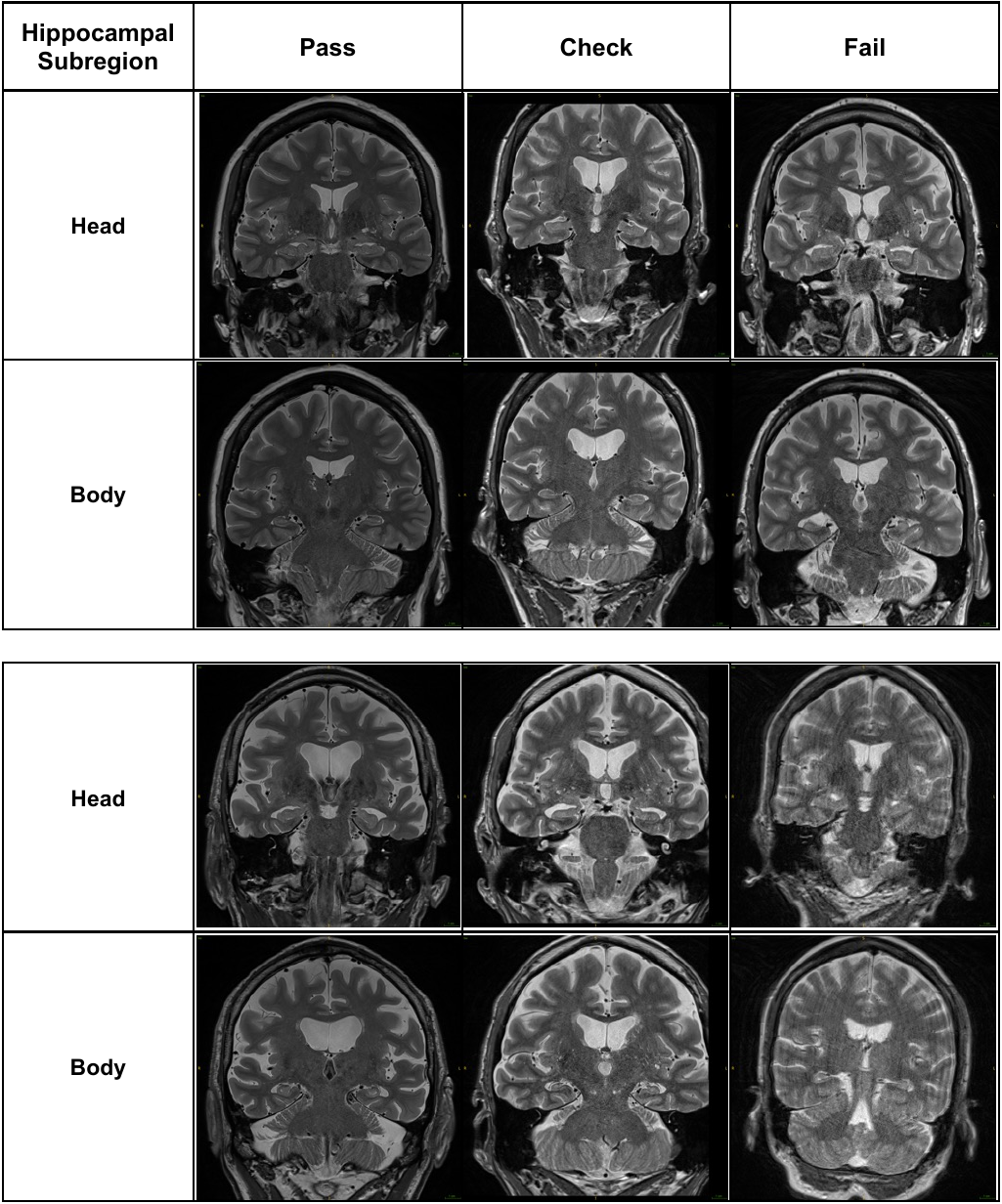

**Figure S1.** Example Pass/Check/Fail of overall image quality ratings from the QC protocol used in Wuestefeld et al. (2023): https://doi.org/10.1093/brain/awad135

**Additional Examples of Landmark QC**

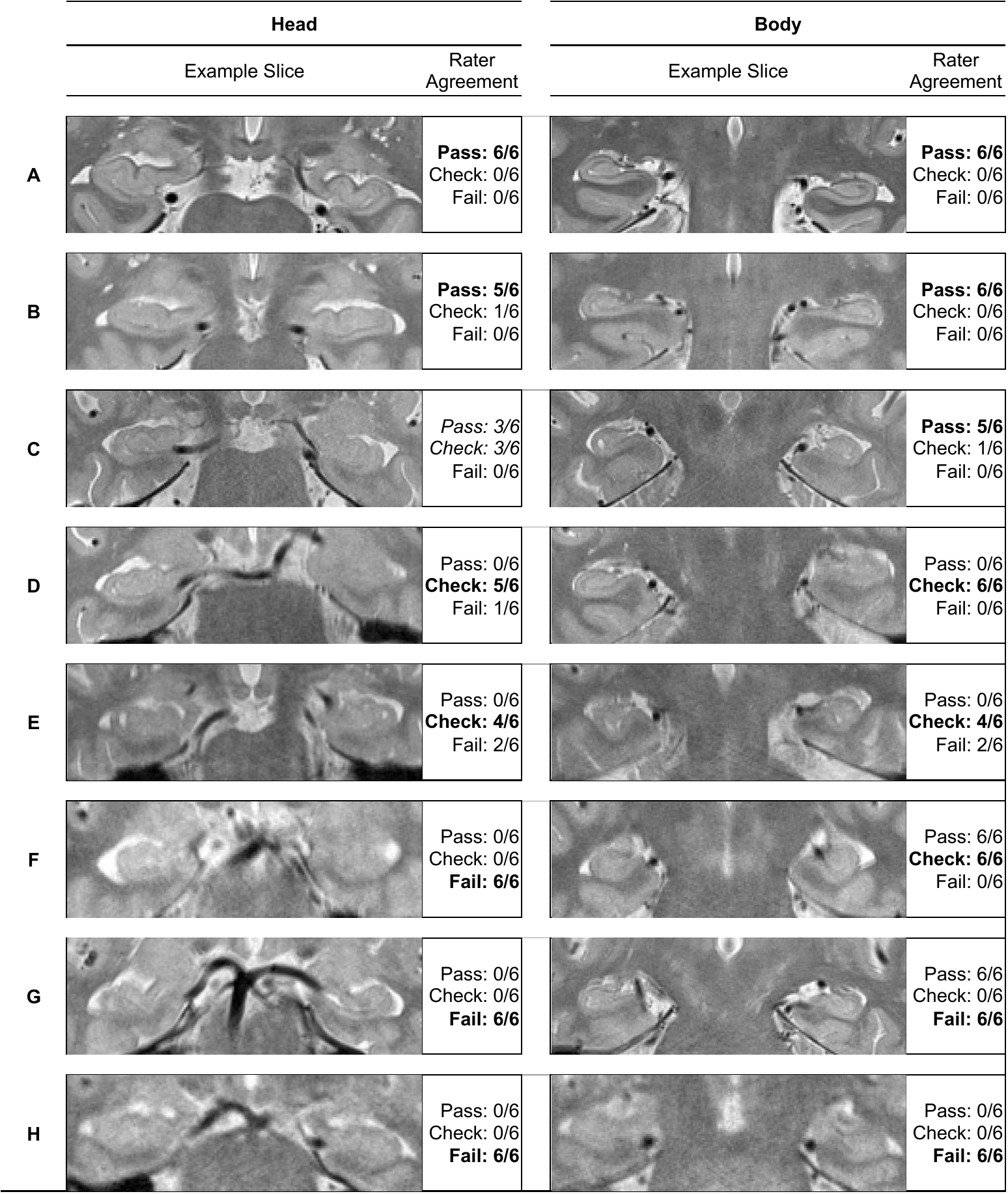

**Figure S2.** Example QC application of a Pass/Check/Fail scale rating landmark visibility required for hippocampal subfield segmentation in head (left panel) and body (right panel). A single rating was determined inclusive of both hemispheres in each hippocampal subregion. Ratings were provided by 6 independent expert raters on 8 individuals (A-H). Scans were collected on the same magnet using the parameters reported in Daugherty et al. (2016). Raters’ decisions were highly consistent, with majority agreement for all but one case (C, Head). Critically, raters unanimously agreed on “Fail” cases, indicating consistent rejection of MR images for automated hippocampal subfield segmentation.

| **Table S4.** Collated qualitative descriptions of process used to determine QC rating by 6 independent experts | |
| --- | --- |
| Prompt | Comments |
| What mental process and/or features of the image did you utilize to reach your final ratings? | - Whether the image was a possible candidate for manual segmentation. - Length of time needed to place the SRLM and/or determine the outer or inner boundary lines. Quick = good; Longer, but possible = fair; No idea = poor. - To be as confident as possible that it is realistic of the subject’s hippocampus; exclude based on judgment of any one hemisphere (lowest rating applied) - Clarity of SRLM, on how many slices and "loose average" of clarity based on majority slices. - Are key landmarks present for each slice; could I trace this slice confidently? All slices = great; 2 slices questionable = fair; more than 2 slices questionable = poor. If guessing for more than 2 slices probably should not be tracing it. - Used experience (compared to past images, does this image seem difficult to trace) |
| Landmarks used to determine | - SRLM - External digitations - Medial edge - Alveus/fimbria |
| Potential steps/adjustments you might take to determine ratings (i.e., between Pass/Check or Check/Fail) | - Apply auto-contrast to the image; but could tweak. - May adjust image contrast on a slice-by-slice value - Scrolling anterior-posterior back and forth to compare slices and visibility. - Zooming in and out to help determine visibility of landmarks. |

**Respondent Provided Article References**

| **Table S5.** Publication examples provided by respondents in which QC methods were used. | |
| --- | --- |
| **#** | **Citation** |
| 1 | ‌ Bender, A. R., Keresztes, A., Bodammer, N. C., Shing, Y. L., Werkle‐Bergner, M., Daugherty, A. M., Yu, Q., Kühn, S., Lindenberger, U., & Raz, N. (2017). Optimization and validation of automated hippocampal subfield segmentation across the lifespan. *Human Brain Mapping*, *39*(2), 916–931. https://doi.org/10.1002/hbm.23891 |
| 2 | Brandão, P. R. de P. (2021). Comprometimento cognitivo na doença de Parkinson : correlatos clínicos, neuropsicológicos e de neuroimagem. *Repositorio.unb.br*. https://repositorio.unb.br/handle/10482/43017 |
| 3 | Canada, K. L., Hancock, G. R., & Riggins, T. (2021). Modeling longitudinal changes in hippocampal subfields and relations with memory from early- to mid-childhood. *Developmental Cognitive Neuroscience*, *48*, 100947. https://doi.org/10.1016/j.dcn.2021.100947 |
| 4 | Canada, K. L., S Saifullah, Gardner, J. C., Sutton, B. P., Fabiani, M., Gratton, G., Raz, N., & Daugherty, A. M. (2023). Development and validation of a quality control procedure for automatic segmentation of hippocampal subfields. *Hippocampus*, *33*(9), 1048–1057. https://doi.org/10.1002/hipo.23552 |
| 5 | Daugherty, A. M., Bender, A. R., Raz, N., & Ofen, N. (2015). Age differences in hippocampal subfield volumes from childhood to late adulthood. *Hippocampus*, *26*(2), 220–228. https://doi.org/10.1002/hipo.22517‌ |
| 6 | DeKraker, J., Ferko, K., Lau, J. C., Köhler, S., & Khan, A. R. (2018). *Unfolding the hippocampus: An intrinsic coordinate system for subfield segmentations and quantitative mapping*. *167*, 408–418. https://doi.org/10.1016/j.neuroimage.2017.11.054‌ |
| 7 | DeKraker, J., Haast, R. A. M., Yousif, M. D., Karat, B., Lau, J. C., Köhler, S., & Khan, A. R. (2022). Automated hippocampal unfolding for morphometry and subfield segmentation with HippUnfold. *ELife*, *11*. https://doi.org/10.7554/elife.77945‌ |
| 8 | DeKraker, J., Lau, J. C., Ferko, K. M., Khan, A. R., & Köhler, S. (2020). Hippocampal subfields revealed through unfolding and unsupervised clustering of laminar and morphological features in 3D BigBrain. *NeuroImage*, *206*, 116328. https://doi.org/10.1016/j.neuroimage.2019.116328‌ |
| 9 | Gervais, N. J., Gravelsins, L., Brown, A., Reuben, R., Karkaby, L., Baker-Sullivan, E., Mendoza, L., Lauzon, C., Almey, A., Foulkes, W. D., Bernardini, M. Q., Jacobson, M., Velsher, L., Rajah, M. N., Olsen, R. K., Grady, C., & Einstein, G. (2022). Scene memory and hippocampal volume in middle-aged women with early hormone loss. *Neurobiology of Aging*, *117*, 97–106. https://doi.org/10.1016/j.neurobiolaging.2022.05.003‌ |
| 10 | Homayouni, R., Yu, Q., Ramesh, S., Tang, L., Daugherty, A. M., & Noa Ofen. (2021). Test–retest reliability of hippocampal subfield volumes in a developmental sample: Implications for longitudinal developmental studies. *Journal of Neuroscience Research*, *99*(10), 2327–2339. https://doi.org/10.1002/jnr.24831‌ |
| 11 | Huguet, J., Falcon, C., Fusté, D., Girona, S., Vicente, D., Molinuevo, J. L., Gispert, J. D., & Operto, G. (2021). Management and Quality Control of Large Neuroimaging Datasets: Developments From the Barcelonaβeta Brain Research Center. *Frontiers in Neuroscience*, *15*. https://doi.org/10.3389/fnins.2021.633438‌ |
| 12 | Keresztes, A., Bender, A. R., Bodammer, N. C., Lindenberger, U., Shing, Y. L., & Werkle-Bergner, M. (2017). Hippocampal maturity promotes memory distinctiveness in childhood and adolescence. *Proceedings of the National Academy of Sciences*, *114*(34), 9212–9217. https://doi.org/10.1073/pnas.1710654114‌ |
| 13 | Keresztes, A., Raffington, L., Bender, A. R., Bögl, K., Heim, C., & Shing, Y. L. (2020). Hair cortisol concentrations are associated with hippocampal subregional volumes in children. *Scientific Reports*, *10*(1). https://doi.org/10.1038/s41598-020-61131-x |
| 14 | Keresztes, A., Raffington, L., Bender, A. R., Bögl, K., Heim, C., & Shing, Y. L. (2022). Longitudinal Developmental Trajectories Do Not Follow Cross-Sectional Age Associations in Hippocampal Subfield and Memory Development. *Developmental Cognitive Neuroscience*, 101085. https://doi.org/10.1016/j.dcn.2022.101085 |
| 15 | Nauer, R. K., Dunne, M. F., Stern, C. E., Storer, T. W., & Schon, K. (2020). Improving fitness increases dentate gyrus/CA3 volume in the hippocampal head and enhances memory in young adults. *Hippocampus*, *30*(5), 488–504. https://doi.org/10.1002/hipo.23166 |
| 16 | Riggins, T., Geng, F., Botdorf, M., Canada, K., Cox, L., & Hancock, G. R. (2018). Protracted hippocampal development is associated with age-related improvements in memory during early childhood. *NeuroImage*, *174*, 127–137. https://doi.org/10.1016/j.neuroimage.2018.03.009 |
| 17 | Santini, T., Koo, M., Farhat, N., Campos, V. P., Alkhateeb, S., Vieira, M. A. C., Butters, M. A., Rosano, C., Aizenstein, H. J., Mettenburg, J., Novelli, E. M., & Ibrahim, T. S. (2021). Analysis of hippocampal subfields in sickle cell disease using ultrahigh field MRI. *NeuroImage: Clinical*, *30*, 102655. https://doi.org/10.1016/j.nicl.2021.102655‌ |
| 18 | Shaw, T., Bollmann, S., Atcheson, N., Strike, L. T., Guo, C., McMahon, K. L., Fripp, J., Wright, M. J., Salvado, O., & Barth, M. (2019). Non-linear realignment improves hippocampus subfield segmentation reliability. *NeuroImage*, *203*, 116206–116206. https://doi.org/10.1016/j.neuroimage.2019.116206‌ |
| 19 | Shaw, T., York, A., Ziaei, M., Barth, M., & Bollmann, S. (2020). Longitudinal Automatic Segmentation of Hippocampal Subfields (LASHiS) using multi-contrast MRI. *NeuroImage*, *218*, 116798. https://doi.org/10.1016/j.neuroimage.2020.116798‌ |
| 20 | Taylor, C. M., Pritschet, L., Olsen, R. K., Layher, E., Santander, T., Grafton, S. T., & Jacobs, E. G. (2020). Progesterone shapes medial temporal lobe volume across the human menstrual cycle. *NeuroImage*, *220*, 117125. https://doi.org/10.1016/j.neuroimage.2020.117125‌ |
| 21 | Wang, L., Miller, J. P., Gado, M. H., McKeel, D. W., Rothermich, M., Miller, M. I., Morris, J. C., & Csernansky, J. G. (2006). Abnormalities of hippocampal surface structure in very mild dementia of the Alzheimer type. *NeuroImage*, *30*(1), 52–60. https://doi.org/10.1016/j.neuroimage.2005.09.017‌ |
| 22 | Wisse, L. E. M., Ungrady, M. B., Ittyerah, R., Lim, S. A., Yushkevich, P. A., Wolk, D. A., Irwin, D. J., Das, S. R., & Grossman, M. (2021). Cross-sectional and longitudinal medial temporal lobe subregional atrophy patterns in semantic variant primary progressive aphasia. *Neurobiology of Aging*, *98*, 231–241. https://doi.org/10.1016/j.neurobiolaging.2020.11.012‌ |
| 23 | Wisse, L. E., Xie, L., Das, S. R., de Flores, R., Hansson, O., Habes, M., Doshi, J., Davatzikos, C., Yushkevich, P. A., & Wolk, D. A. (2022). Tau pathology mediates age effects on medial temporal lobe structure. *Neurobiology of Aging*, *109*, 135–144. https://doi.org/10.1016/j.neurobiolaging.2021.09.017 |
| 24 | Wisse, L., Robin de Flores, Xie, L., Das, S. R., McMillan, C. T., Trojanowski, J. Q., Grossman, M., Lee, E. B., Irwin, D. J., Yushkevich, P. A., & Wolk, D. A. (2021). Pathological drivers of neurodegeneration in suspected non-Alzheimer’s disease pathophysiology. *Alzheimer’s Research & Therapy*, *13*(1). <https://doi.org/10.1186/s13195-021-00835-2> |
| 25 | Woodworth, D. C., Nguyen, H. L., Khan, Z., Kawas, C. H., Corrada, M. M., & S. Ahmad Sajjadi. (2020). Utility of MRI in the identification of hippocampal sclerosis of aging. *Alzheimer’s & Dementia*, *17*(5), 847–855. https://doi.org/10.1002/alz.12241‌ |
| 26 | Woodworth, D. C., Sheikh-Bahaei, N., Scambray, K. A., Phelan, M. J., Perez-Rosendahl, M., Corrada, M. M., Kawas, C. H., & Sajjadi, S. A. (2022). Dementia is associated with medial temporal atrophy even after accounting for neuropathologies. *Brain Communications*, *4*(2). https://doi.org/10.1093/braincomms/fcac052‌ |
| 27 | Xie, L., Shinohara, R. T., Ittyerah, R., Kuijf, H. J., Pluta, J. B., Blom, K., Kooistra, M., Reijmer, Y. D., Koek, H. L., Zwanenburg, J. J. M., Wang, H., Luijten, P. R., Geerlings, M. I., Das, S. R., Biessels, G. J., Wolk, D. A., Yushkevich, P. A., & Wisse, L. E. M. (2018). Automated Multi-Atlas Segmentation of Hippocampal and Extrahippocampal Subregions in Alzheimer’s Disease at 3T and 7T: What Atlas Composition Works Best? *Journal of Alzheimer’s Disease*, *63*(1), 217–225. https://doi.org/10.3233/JAD-170932 |
